# A fungal ribonuclease-like effector protein inhibits plant host ribosomal RNA degradation

**DOI:** 10.1101/291427

**Authors:** Helen G. Pennington, Rhian Jones, Seomun Kwon, Giulia Bonciani, Hannah Thieron, Thomas Chandler, Peggy Luong, Sian Morgan, Michal Przydacz, Tolga Bozkurt, Sarah Bowden, Melanie Craze, Emma Wallington, James Garnett, Mark Kwaaitaal, Ralph Panstruga, Ernesto Cota, Pietro D. Spanu

## Abstract

The biotrophic fungal pathogen *Blumeria graminis* causes the powdery mildew disease of cereals and grasses. Proteins with a predicted ribonuclease (RNase)-like fold (termed RALPHs) comprise the largest set of secreted effector candidates within the *B. graminis* f. sp. *hordei* genome. Their exceptional abundance suggests they play crucial functions during pathogenesis. We show that transgenic expression of RALPH CSEP0064/BEC1054 increases susceptibility to infection in monocotyledenous and dicotyledonous plants. CSEP0064/BEC1054 interacts *in planta* with five host proteins: two translation elongation factors (eEF1α and eEF1γ), two pathogenesis-related proteins (PR5 and PR10) and a glutathione-S-transferase. We present the first crystal structure of a RALPH, CSEP0064/BEC1054, demonstrating it has an RNase-like fold. The protein interacts with total RNA and weakly with DNA. Methyl jasmonate levels modulate susceptibility to aniline-induced host RNA fragmentation. *In planta* expression of CSEP0064/BEC1054 reduces the formation of this RNA fragment. We propose that CSEP0064/BEC1054 is a pseudoenzyme that binds to host ribosomes, thereby inhibiting the action of plant ribosome-inactivating proteins that would otherwise lead to host cell death, an unviable interaction and demise of the fungus.

## INTRODUCTION

The obligate biotrophic fungus *Blumeria graminis* causes powdery mildew disease on some small grain cereals and grasses (Poaceae). A high degree of host specificity is displayed by the fungus, with at least eight *formae speciales* (f.sp.), each infecting a different host genus. *Blumeria graminis* f.sp. *hordei* and *B. graminis* f.sp. *tritici* colonise barley (*Hordeum vulgare* L.) and wheat (*Triticum aestivum* L.), respectively, and they can result in large crop losses (Troch et al., 2014). In the case of powdery mildews, the only part of the fungus that actually penetrates the plant during infection is the haustorium (O’Connell & Panstruga, 2006), a dedicated fungal infection structure thought to absorb plant nutrients. The success of infection is determined by the outcome of a “secretory warfare” between the host and the pathogen, which produces effectors at the haustorial complex (O’Connell & Panstruga, 2006, Panstruga & Dodds, 2009).

Genomes of powdery mildew fungi code for several hundred effector-like proteins: 491 Candidate Secreted Effector Proteins (CSEPs) were identified in the *B. graminis* f.sp. *hordei* genome (Pedersen et al., 2012) and 437 in the *B. graminis* f.sp. *tritici* genome (Wicker et al., 2013). These putative effectors have a predicted amino-terminal signal peptide for secretion, lack recognizable transmembrane domains, and have no relevant BLAST hits outside of the powdery mildew family (Erysiphaceae). As for most other pathogen effectors, their mode of action is not yet understood. Previously, two complementary bioinformatic procedures were used to identify CSEPs with predicted RNA-binding or ribonuclease folds: InterProScan combined with Gene ontology (GO) characterization, and IntFOLD, an integrated structure prediction server. In total, 72 out of the 491 *B. graminis* f.sp. *hordei* CSEPs were found by these approaches, with 54 predicted by InterProScan and 37 by IntFOLD (19 of which were found by both techniques (Pedersen et al., 2012)).

Proteins occurring within CSEP families with structural similarity to ribonuclease and/or RNA-binding activity were termed RNase Like Proteins expressed in Haustoria (RALPH)s (Spanu, 2017). Some of the CSEP families contain members for which the RNase domain is not recognised by the prediction algorithms. If the latter are included, the RALPHs comprise the biggest subset of effector candidates within the *Blumeria graminis* f. sp. *hordei* genome, numbering at least 113 of the 491 CSEPs. Their abundance, and their proliferation within a genome that otherwise has lost numerous genes (Spanu et al., 2010), suggests that they play a prominent role during infection (Pedersen et al., 2012). The genes encoding RALPHs, and other CSEPs, are often physically closely linked to (retro-)transposable elements, indicating that the duplication of these effectors may have occurred through illegitimate recombination of (retro-)transposon sequences (Pedersen et al., 2012). An RNA interference (RNAi)-based screen for functionally important effector genes in *B. graminis* f.sp. *hordei* identified eight genes whose expression is required for full pathogenic development. These included two genes encoding RALPH effectors: CSEP0064/BEC1054 and CSEP0264/BEC1011 (Pliego et al., 2013).

We have previously found four barley proteins that interact with CSEP0064/BEC1054 using *in vitro* affinity assays followed by liquid chromatography mass spectrometry (LCMS) analysis. These proteins comprise a eukaryotic Elongation Factor 1 gamma (eEF1γ), a Pathogenesis Related Protein 5 (PR5), a Glutathione-S-Transferase (GST) and a Malate Dehydrogenase (MDH). The respective protein-protein interactions were confirmed by subsequent yeast two-hybrid (Y2H) experiments (Pennington et al., 2016a). Other barley proteins bind to CSEP0064/BEC1054 *in vitro* but do not interact in yeast. These are PR10, a ribosomal 40S subunit protein 16 (40S 16), and a eukaryotic Elongation Factor 1 alpha (eEF1α). Whilst affinity-LCMS and Y2H approaches possess many strengths, both techniques are associated with the identification of false positives, for example, proteins that bind the affinity matrix in LCMS, or proteins that interfere with the reporter readouts in Y2H (Brückner et al., 2009, Płociński et al., 2014). Alternative methods can be used to address these concerns, for example *in planta* Bimolecular Fluorescence Complementation (BiFC; (Ghosh et al., 2000)) assays.

The predicted ribonuclease structure of CSEP0064/BEC1054 was originally determined by *in silico* homology modelling. However, the protein, like other RALPHs, is unlikely to be catalytically active, because it lacks the active site residues known to be required for RNase activity (Pedersen et al., 2012). Plants also possess RNases that are proposed to be implicated in defence against pathogens. Examples are the ribosome-inactivating proteins (RIPs), which depurinate the sarcin-ricin loop (SRL) in the ribosomal RNA (rRNA) of the large ribosome subunit (Reinbothe et al., 1994b). In barley, the jasmonate-induced protein of 60 kDa (JIP60) is a RIP involved in mediating host-induced cell death (Chaudhry et al., 1994, Reinbothe et al., 1994a).

Here, we show that heterologous transgenic expression of CSEP0064/BEC1054 in wheat enhances susceptibility to *B. graminis* f.sp. *tritici.* Similarly, expression of this protein in *Nicotiana benthamiana* increases the susceptibility to the oomycete pathogen *Peronospora tabacina.* To provide mechanistic insights into the function of CSEP0064/BEC1054, we solved the structure of the first RALPH by X-ray crystallography and studied the protein by nuclear magnetic resonance (NMR) spectroscopy, determining unequivocally a high degree of structural similarity with fungal RNases. Given the similarity of the predicted structure of *Blumeria* RALPH proteins to fungal RIPs, we hypothesise that RALPHs may stoichiometrically outcompete host RIPs, thereby inhibiting their cell death-promoting activity and thus act as a pseudoenzyme (Eyers & Murphy, 2016). We present evidence that CSEP0064/BEC1054 interferes with methyl-jasmonate induced cleavage of RNA in wheat.

## RESULTS

### CSEP0064/BEC1054 increases susceptibility to adapted pathogens

We previously assessed the contribution of CSEP0064/BEC1054 to the interaction of barley and its adapted powdery mildew pathogen, *B. graminis* f.sp. *hordei*, by Host-Induced Gene Silencing (HIGS; (Pliego et al., 2013)). In this native context, CSEP0064/BEC1054 contributes significantly to fungal virulence: when the respective gene is silenced by HIGS, the infection success (haustorial index) drops to less than half of the value obtained with a control construct.

To explore the effect of CSEP0064/BEC1054 on interactions with additional pathogens, we generated transgenic bread wheat (*T. aestivum*) lines constitutively expressing a codon-modified version of *CSEP0064/BEC1054* lacking the N-terminal signal peptide for secretion (termed *wBEC1054*), which was also used in our previous HIGS experiments (Pliego et al., 2013). We selected progeny of three homozygous T3 lines, two carrying single expressed copies of the *CSEP0064/BEC1054* transgene (+/+, lines 3.3.7 and 3.3.14) and one null segregant (lacking the transgene), referred to as azygous (-/-, line 3.3.12), which served as a negative control in our study.

We first determined whether expression or potential unintended gene disruption by the transgene affects morphology and/or development of the wheat lines. To this end, we investigated the phenotype of adult T4 plants homozygous for the effector transgene (+/+) or respective azygous controls (-/-) under the same experimental conditions as used for subsequent assays. According to common practice for the phenotypic characterization of wheat plants (Borrell et al., 1993, Fellahi et al., 2013, Gasperini et al., 2012, Hays et al., 2007), eleven characteristics were assayed quantitatively: leaf number, maximum height, peduncle (internode 1) and other internode lengths, ear length, subcrown length, fertile tiller number, tiller mass and grain number. The respective values for these parameters from the azygous individuals and the transgenics were indistinguishable (Supplemental Figure 1), that is the presence of transgenic *CSEP0064/BEC1054* does not affect the adult phenotype of wheat, in our experimental conditions.

We next measured the effect of the *CSEP0064/BEC1054* transgene on the susceptibility of the wheat lines to its adapted powdery mildew pathogen, *B. graminis*f.sp. *tritici*. Microcolonies with epiphytic hyphae formed can be used as a proxy for the presence of functional haustoria. This parameter can be expressed as the proportion of microcolonies relative to the number of germinated conidia (propH; Figure 1A). We determined the propH value at the base, the middle and the tip of both young and mature leaves of T4 individuals of the transgenic lines (+/+) and the azygous plants (-/-). The median propH was consistently higher in leaf blades of plants derived from the transgenic lines (+/+) as compared to the controls (-/-), irrespective of the position within the leaf (base, middle, tip) or the age of the leaf (young, mature; Figure 1A). Notably, the spread of the data increased from the leaf base to the leaf tip in both young and mature leaves, as shown by the increasing size of the boxes in the boxplots and their respective error bars (Figure 1A).

**Figure 1.**
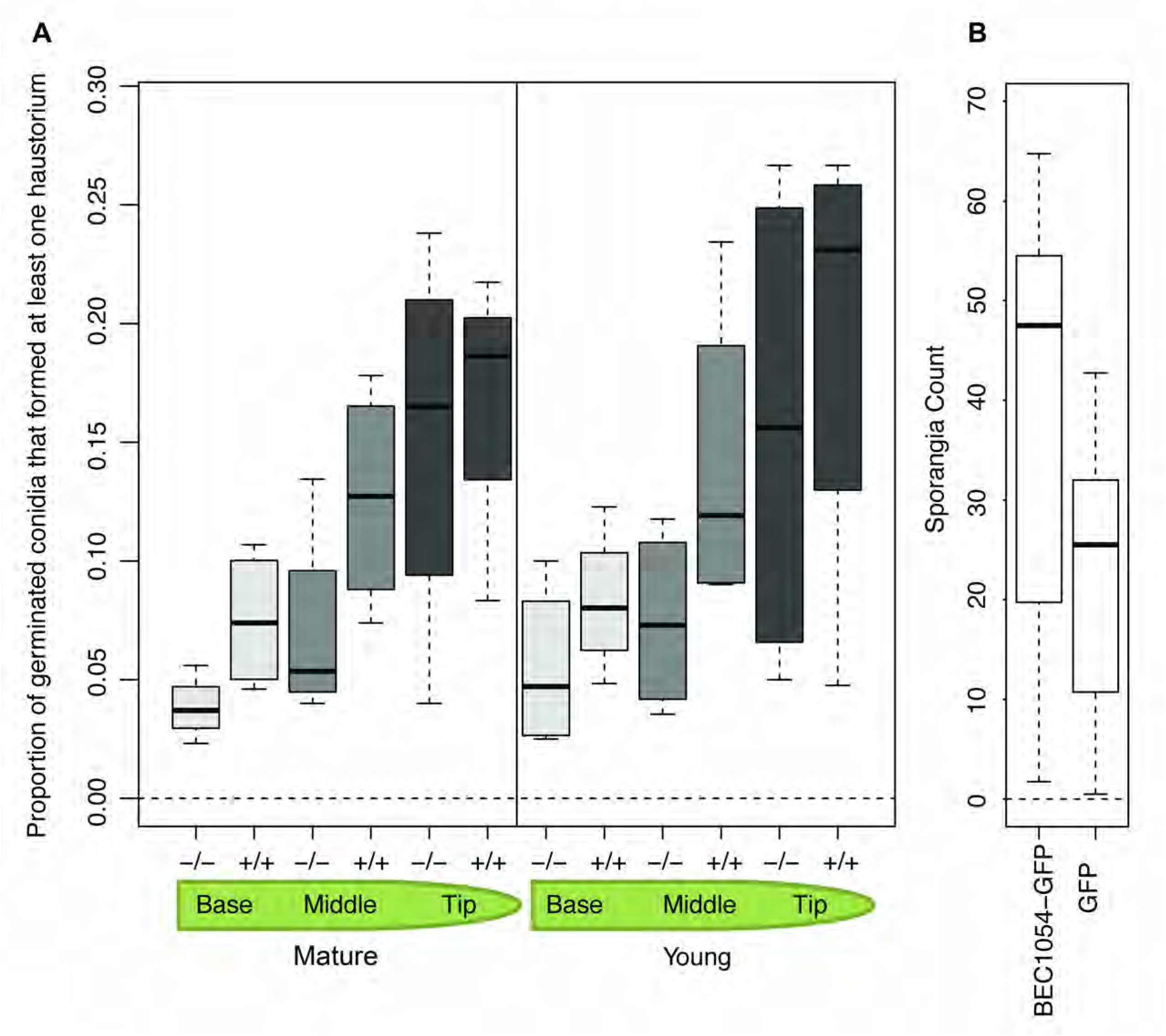
Transgenic expression of CSEP0064/BEC1054 in plants enhances susceptibility to adapted pathogens. **A**) The presence of the transgene *wbec1054,* encoding CSEP0064/BEC1054 from the non-adapted barley powdery mildew pathogen, increases haustorium formation of the adapted pathogen *B. graminis* f.sp. *tritici* in wheat. Two centimetre leaf segments were taken from primary leaves (young plants), or from the third most recent leaf (mature plants), and the mean proportion of germinated conidia that produced a functional haustorium (determined by measuring the number of colonies forming an extensive network of epiphytic hyphae) was assessed. Plants were either homozygous (+/+) or azygous (-/-) for *wbec1054*. Young plants were three weeks old, and mature plants were 11 weeks old. The boxes represent the quartiles, the thick line denotes the median, and maximum and minimum values are shown by the error bars. **B**) Sporangia production by the adapted downy mildew pathogen *P. tabacina* is increased in *N. benthamiana* by expression of CSEP0064/BEC1054 from the non-adapted barley powdery mildew pathogen. *N. benthamiana* leaves were infiltrated on one side of the midrib with Agrobacteria expressing GFP and RFP, and on the other side with Agrobacteria expressing CSEP0064/BEC1054 with a C-terminal GFP tag and RFP (as a transformation marker). Leaves were inoculated, within one hour of infiltration, with *P. tabacina* sporangia on their abaxial surface. After 10 days, leaf disks were collected from each leaf, the sporangia removed via washing, counted, and used to calculate the mean. Significantly more sporangia were found with CSEP0064/BEC1054-GFP than with GFP only (n=5, p<0.001). The thick line denotes the median of each boxplot, the boxes represent the quartiles, maximum and minimum values are shown by the error bars.

To test the contribution of CSEP0064/BEC1054 to a plant-microbe interaction with a dicotyledonous host species and an adapted pathogen different from powdery mildew, we opted for transient expression of *CSEP0064/BEC1054* in *Nicotiana benthamiana* and subsequent challenge with the oomycete pathogen *Phytophthora tabacina*, the causal agent of the tobacco downy mildew disease. *Agrobacterium tumefaciens*-mediated transformation was used to transiently express either Green Fluorescent Protein (GFP)-tagged CSEP0064/BEC1054 (C terminal tag), or GFP alone as a control, in *N. benthamiana* (in each case co-expressed with RFP as a transformation marker). Leaves from four-week-old leaves were detached, and co-infiltrated with *Agrobacterium* expressing CSEP0064/BEC1054-GFP and RFP on one side of the midrib, and with *Agrobacterium* co-expressing GFP and RFP on the other. The leaves were then inoculated with *P. tabacina* sporangia on the abaxial surface, and the number of sporangia produced on the leaf after 10 days was assayed. The presence of GFP-tagged CSEP0064/BEC1054 significantly increased the mean of the sporangia produced as compared to the GFP control (Figure 1B).

In conclusion, transgenic expression of CSEP0064/BEC1054 promotes virulence of diverse adapted pathogens (*B. graminis* f.sp. *tritici*, *P. tabacina*) in monocotyledonous (wheat) and dicotyledonous (*N. benthamiana*) plant species.

### *In planta* validation of CSEP0064/BEC1054 protein interactors

In order to validate physical *in planta* interactions between CSEP0064/BEC1054 and several host proteins previously identified by protein *in vitro* affinity-LCMS experiments and targeted Y2H assays (Pennington et al., 2016a), we performed Bimolecular Fluorescence Complementation (BiFC) experiments with the fungal effector and the respective candidate targets (Hu et al., 2002). Besides the CSEP0064/BEC1054 bait, we used the RIP JIP60ml, a processed and constitutively active form of barley JIP60 (Rustgi et al., 2014), as a (negative) control bait in our experiments, as we previously found no detectable direct interactions between JIP60ml and the barley prey proteins *in vitro* (Pennington et al., 2016a).

We first analysed the *in planta* subcellular localization of all putative interaction partners (two bait and nine barley prey proteins). The bait and prey proteins were translationally fused at the C-terminus with monomeric yellow fluorescent protein mYFP) and transiently co-expressed with RFP (as a transformation marker) in barley leaf epidermal cells (Panstruga, 2004). The exception was PR5, which was fused to mYFP at the N-terminus in order to mask the signal peptide encoded by the gene and to prevent secretion. Expression and subcellular localization were determined via confocal microscopy (Figure 2A). In the case of CSEP0064/BEC1054-mYFP and JIP60ml-mYFP, there was a weak and diffuse fluorescence signal in the cytoplasm and the nucleus; in addition, a punctate distribution of both proteins was observed in the cytoplasm. The mYFP fusion proteins of PR5 and PR10 were distributed fairly homogeneously throughout the cytoplasm, with brighter fluorescence evident in the nucleus. The GST-mYFP fusion protein yielded a very diffuse and faint signal in the cytoplasm and nucleus, with bright puncta in the cytoplasm, and a bright signal in a section of the nucleus. Fluorescence of three elongation factor fusions (eEF1γ-mYFP, eEF1α(1)-mYFP and eEF1α(3)-mYFP) was detectable in the cytoplasm, and for eEF1α(1)-mYFP and eEF1α(3)-mYFP (two paralogs of eEFA we cloned from barley) fluorescence was also observable in the nucleus. The 40S S16-mYFP fusion protein yielded a very diffuse and faint signal in the cytoplasm and the nucleus, with bright puncta in the cytoplasm, and a bright signal in the nucleus. The NDPK-(used as a negative control in previous Y2H assays; (Pennington et al., 2016a)) and MDH-mYFP fusions were both observed in the nucleus and cytoplasm (Figure 2A). Taken together, all bait and prey fusion proteins yielded detectable *in planta* fluorescence and were present in the cytoplasm and in part in the nucleus, allowing for potential interaction in these cellular compartments.

**Figure 2.**
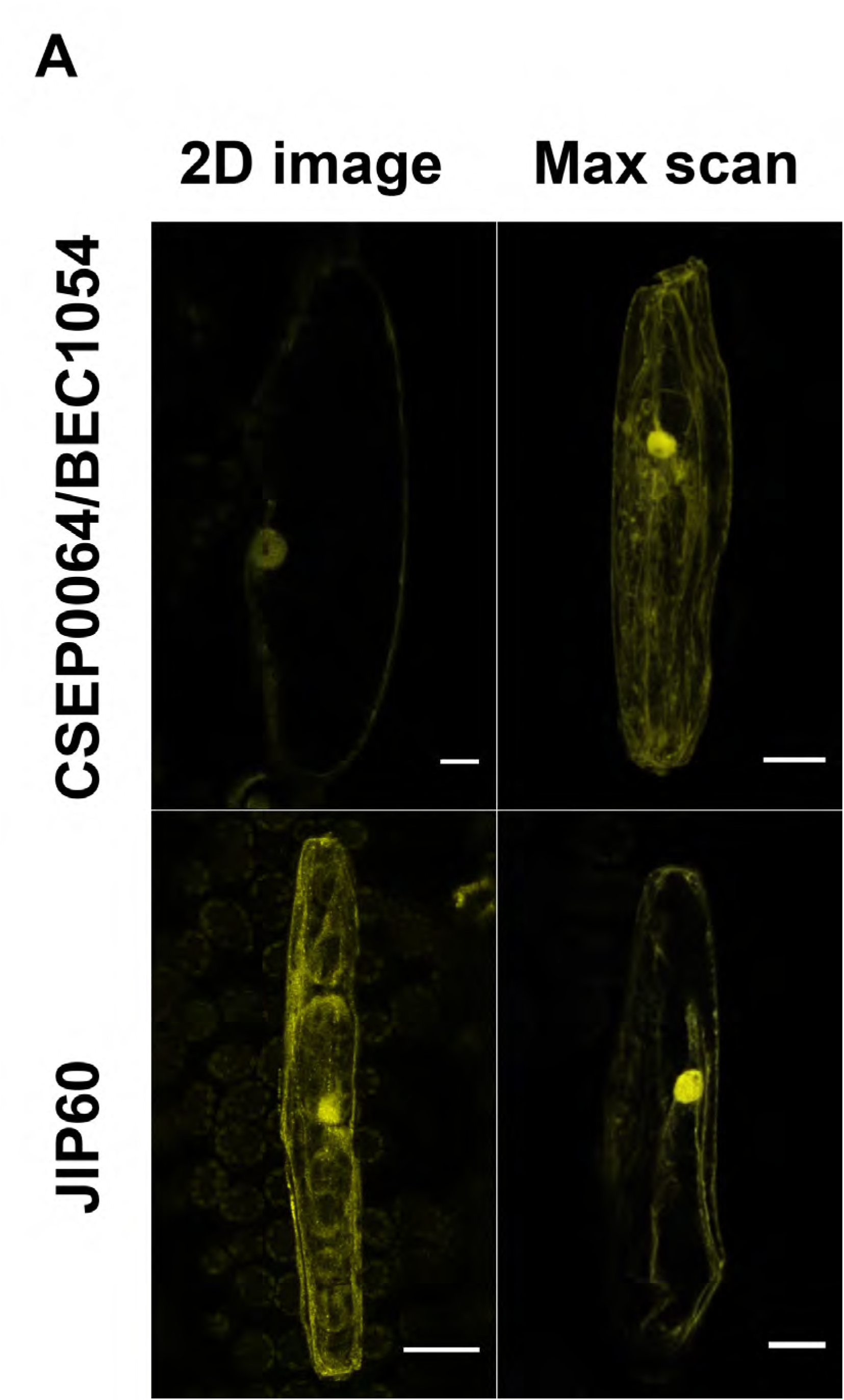
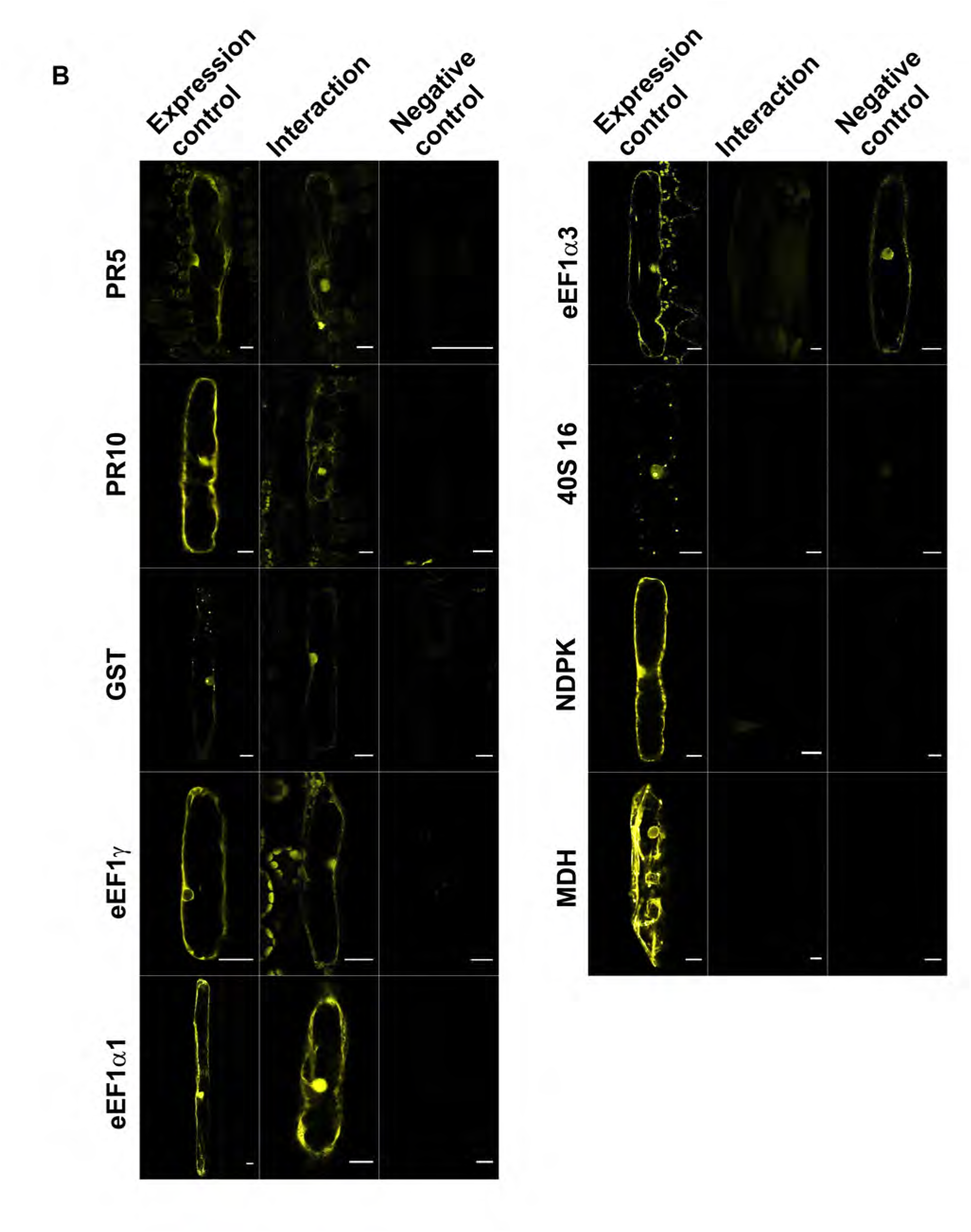
CSEP0064/BEC1054 interacts with multiple proteins *in planta*. **A**) Micrographs show single barley leaf epidermal cells, transformed by particle bombardment, transiently co-expressing RFP (as a transformation marker) and the mYFP-tagged bait proteins Jasmonate-induced protein 60 (JIP60) and CSEP0064/BEC1054. The “Max scan” was created through combining maximum pixel intensity of Z-stack images (six and 23 images, respectively) whereas the 2D image represents a single confocal image. The JIP60 protein contains an internal peptide, which inhibits its activity, which was removed and replaced with a methionine-leucine linker, and is therefore referred to as “JIP60ml” (see Materials and Methods for details). Scale bars are 20 µM. **B**) The host plant, barley, was used to perform a BiFC assay via particle bombardment-mediated transient gene expression in single leaf epidermal cells. The fluorescent signal of the prey proteins under investigation, tagged with mYFP, is shown in the “Expression control” column. The “Interaction” column shows the fluorescence signal seen upon co-expression of the respective prey protein with CSEP0064/BEC1054 and RFP (as a transformation marker). The “Negative control” column indicates the fluorescence seen if the prey proteins were coexpressed with JIP60. The JIP60 protein contains an internal peptide, which inhibits its activity, which was removed and replaced with a methionine-leucine linker, and is therefore referred to as “JIP60ml” (see Materials and Methods for details). The barley prey proteins were: Pathogenesis-Related protein 5 (PR5), PR10, and Glutathione-S-Transferase (GST), eukaryotic Elongation Factor 1 Gamma (eEF1γ), eukaryotic Elongation Factor 1 Alpha (eEF1α) isoforms 1 and 3, nucleoside diphosphate kinase (NDPK; negative control) and malate dehydrogenase (MDH). Scale bars are 20 µM.

A clear BiFC signal was seen in each case when CSEP0064/BEC1054, C-terminally tagged with the C-terminal half of mYFP, was co-expressed with RFP (as a transformation marker) and the five prey proteins C-terminally tagged with the N-terminal domain of mYFP (except for PR5, which was N-terminally tagged with the N-terminal domain of mYFP): PR5, PR10, GST, eEF1γ and eEF1α(1) (Figure 2B). In case of these bait-prey pairings, mYFP fluorescence was observed predominantly in the nucleus and in most cases also weakly in the cytoplasm. This pattern largely matched the subcellular localisation of the bait protein CSEP0064/BEC1054 expressed alone (Figure 2A). There was no evident mYFP fluorescence when these tagged proteins were co-expressed with the negative control, JIP60ml. We also detected no apparent mYFP signal when the CSEP0064/BEC1054 bait protein was co-expressed with the remaining four prey proteins (eEF1α(3), MDH, 40S S16 and the NDPK negative control; Figure 2B). Likewise, fluorescence was lacking upon co-expression of the JIP60ml bait with MDH, 40S S16 and NDPK; Figure 2B). We observed, however, a signal in the cytoplasm and the nucleus when JIP60ml was co-expressed with eEF1α(3) (Figure 2B).

### CSEP0064/BEC1054 shows structural similarity to T1 RNases

To provide detailed insights into the molecular architecture of RALPH effectors, we determined the structure of CSEP0064/BEC1054 by X-ray crystallography. An initial model obtained from a dataset phased by iodide-SAD (single-wavelength anomalous diffraction) was subsequently used to calculate phases for a native dataset at 1.3 Å via molecular replacement. Data processing and refinement statistics for the structures are outlined in Supplemental Table 1. The crystal structure reveals a canonical (α+β) T1 RNase fold for CSEP0064/BEC1054. This protein comprises a core four-stranded anti-parallel β-sheet packed against an α-helix, and a second shorter peripheral anti-parallel β-sheet formed from the N terminal hairpin and C-terminal strand β7 (Figure 3A). This arrangement is constrained by a disulphide bridge formed between residues C6 and C92, a feature that is universally conserved in the T1 RNase family (Hill et al., 1983).

**Figure 3.**
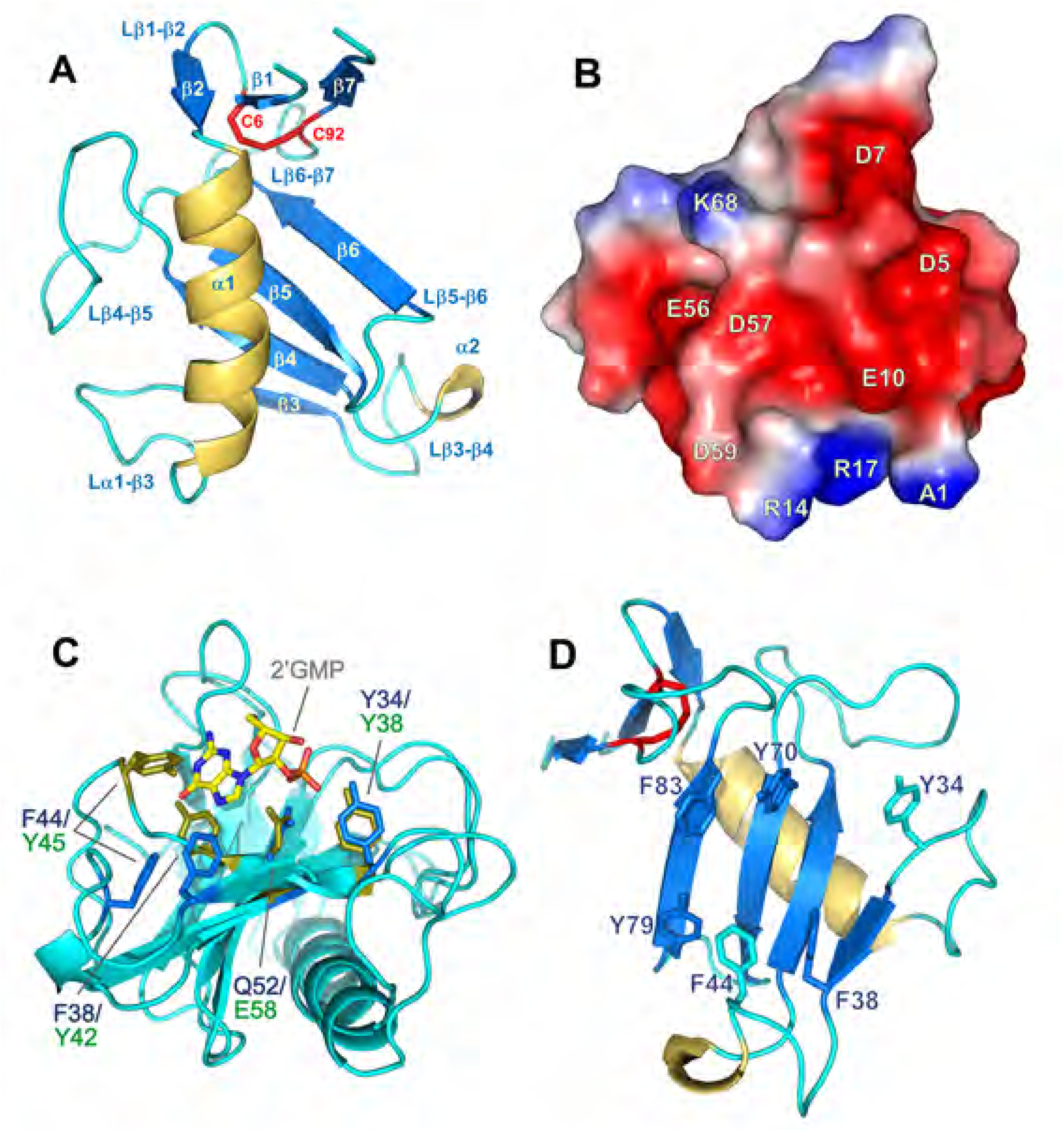
Crystal structure of CSEP0064/BEC1054. **A**) CSEP0064/BEC1054 exhibits a canonical T1 RNase type fold, including a disulphide bridge between C6 and C92 linking the N and C-termini of the protein (shown in red). **B**) Electrostatic surface potential calculated using APBS (Dolinsky et al., 2007), shows a patch (shown in red) of negatively charged residues on the surface of CSEP0064/BEC1054. **C**) Structural overlay of the CSEP0064/BEC1054 structure with F1 RNase of the T1 family (2.5 Å RMSD for Cα atoms of the secondary structural elements). Conserved residues in the putative RNA (2’ guanosine monophosphate) binding site are highlighted in blue and green for CSEP0064/BEC1054 and the F1 RNAse, respectively. **D**) Exposed aromatic side chains in the large β-sheet and the α1-β3, β3-4 loops cluster on the concave face of the CSEP0064/BEC1054 protein.

Submission of the CSEP0064/BEC1054 structural coordinates to the Dali server (Holm & Rosenström, 2010), confirmed that RNase T1 family members are the closest structural homologues of CSEP0064/BEC1054, a family of RNases that show a high degree of specificity for cleavage of substrates at the 3’ end of guanosyl residues (Yoshida, 2001). Charge is clearly segregated at the CSEP0064/BEC1054 surface, as the majority of negatively charged side chains cluster in the vicinity of the small β sheet (strands β1, β2 and β7; residues D5, D7, E10, E13, E56, D57, D59, E81 and E96). This surface is bordered by side chains of R14, R17, K68 and K91, whilst the opposite side of the protein is predominantly cationic (R27, K28, R51, K75, R76; Figure 3B). Interestingly, this feature is not observed in the structures of the prototypical T1 RNase from *Aspergillus oryzae* (Martinez-Oyanedel et al., 1991) or the F1 RNase from *Fusarium moniliforme* (Vassylyev et al., 1993).

Whilst CSEP0064/BEC1054 lacks the catalytic triad (E58, H92, R77) of the T1 RNases (Pedersen et al., 2012), analysis of the structure reveals several similarities at the level of the putative substrate binding site. When overlaid with the structure of T1 RNase, residues located in loop α1-β3 (R27/K28) and at the end of loop β6-β7 (K95) flank the β sheet containing the putative RNA-binding site, and thus could feasibly interact with the anionic phosphate backbone of nucleic acids. Moreover, several residues that interact with RNA substrates in other T1-like structures are conserved in CSEP0064/BEC1054 (Figure 3C). Comparison with the F1 RNase reveals that residues Y34 and Y70 in CSEP0064/BEC1054 can recreate the contacts of residues Y38 and R77 in RNase F1 with the phosphate of 2’ guanosyl monophosphate. Additionally, residues F38 and F44 in CSEP0064/BEC1054 are structurally equivalent to Y42 and Y45 of this RNase, involved in π stacking interactions with the guanosine base (Vassylyev et al., 1993). However, residues N43, N44 and N98, which form hydrogen bonds to the guanosyl base substrates in the F1 RNase, are not conserved in CSEP0064/BEC1054. These residues have been identified as critical to conferring guanosyl binding specificity (Steyaert et al., 1991).

More generally, the concave face of CSEP0064/BEC1054, defined by the main β-sheet and extended loops β3-β4 and β6-7 (Figure 3D), is notably hydrophobic for a solvent-exposed surface and is lined with aromatic residues (Y34, F38, F44, Y70, Y79 and F83). Such aromatics could have a role in base-stacking interactions with nucleic acids, or could constitute a binding site for alternative protein ligands that require a hydrophobic surface. Indeed, hydrophobic surface patches have also been identifed in AVR1-CO39 and AVR-Pia effector proteins from the rice blast fungus *Magnaporthe oryzae*; two of the few structurally characterised fungal effectors to date (de Guillen et al., 2015).

### NMR solution studies agree with the CSEP0064/BEC1054 crystal structure and highlight regions of conformational flexibility

CSEP0064/BEC1054 was further structurally characterised in solution via NMR spectroscopy following isotopic ^15^N and ^13^C labelling of the protein. Figure 4 shows the ^1^H-^15^N HSQC spectrum for CSEP0064/BEC1054, where each crosspeak corresponds to amide groups in the protein. Good crosspeak dispersion in this spectrum indicates that the protein is monomeric and well-folded in solution. Using conventional 3D experiments, 89 out of 94 backbone amides were assigned (i.e. excluding the backbone imines of 3 prolines). Residues that could not be assigned (S42, F44, G46, G64 and S88) display large crystallographic B factors. These residues map to loop regions on opposite faces of the core β sheet, suggesting that they experience motions that prevent sampling of a defined chemical environment (e.g. at intermediate exchange regime in NMR timescales). Intriguingly, comparison with the majority of T1 RNase structures crystallised with ligands typically show these loops packed in a “closed” conformation over the core β sheet, and it is feasible that their flexibility could have a role in substrate binding.

**Figure 4.**
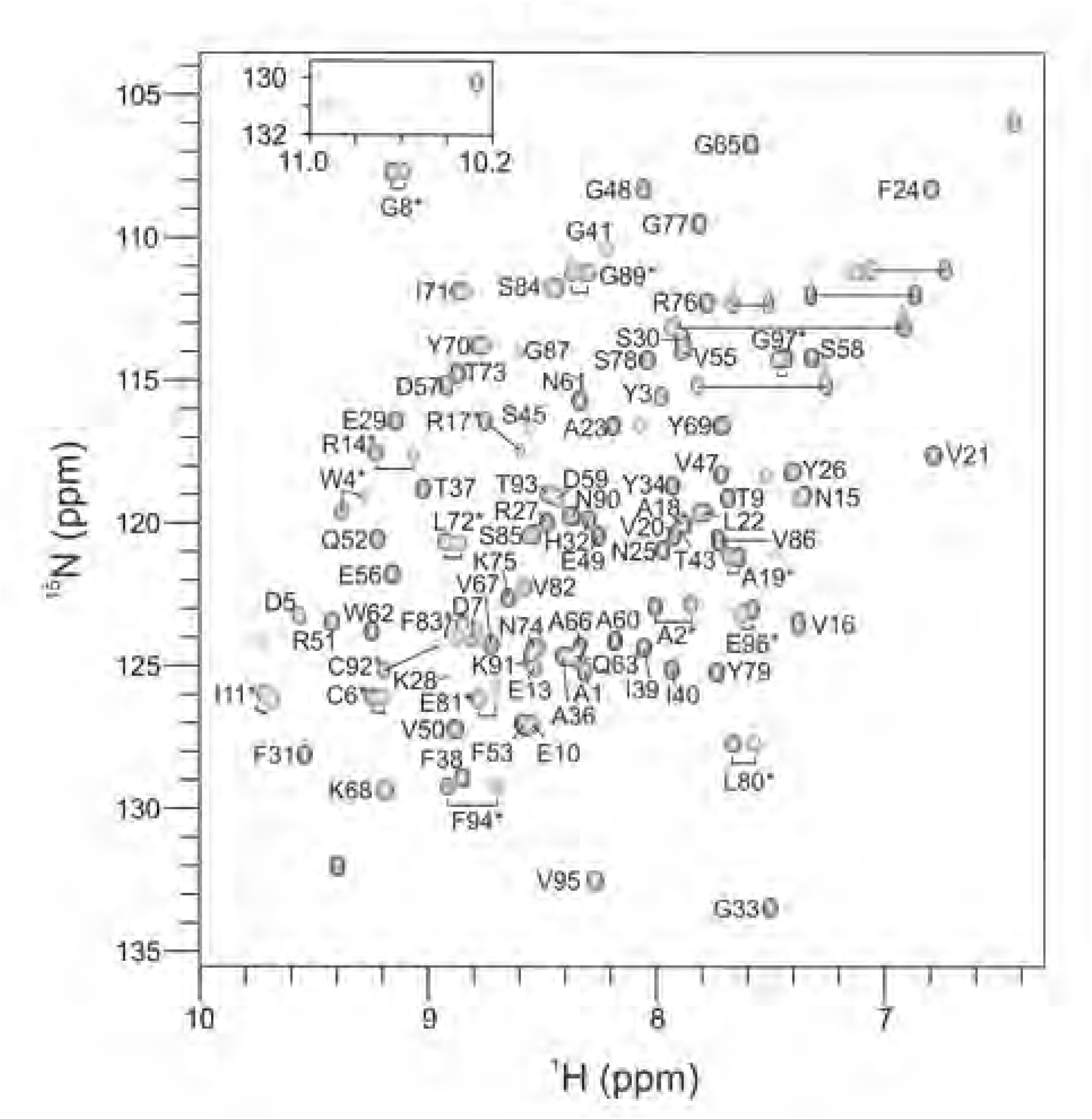
^1^H-^15^N HSQC spectrum showing NMR backbone assignments of CSEP0064/BEC1054. Spectra used to obtain these assignments were recorded in 50 mM sodium phosphate, 150 mM NaCl, pH 7.4 at 308 K and 600 MHz. Residues with doubled amide crosspeaks in the ^1^H and ^15^N frequencies (suggesting alternative conformations due to *cis-trans* isomerization of prolines 12 and 54) are marked with an asterisk. Resonances from side chain amides (upper right) are connected by a straight line.

In addition, we observed doubling of crosspeaks for amides of residues located in strands β1, β6, β7 and the α1 helix (A2, W4, C6, G8, I11, R14, R17, A19, I39, L72, L80, F83, G89, C92, F94, E96 and G97), suggesting that this region adopts two distinct conformations in solution, possibly induced by *cis* and *trans* configurations of neighbouring prolines 12 and 54. Positions of secondary structure elements predicted from NMR chemical shift frequencies by the program TALOS (Shen et al., 2009) concur well with those observed in the crystal structure. All these elements are well structured in solution, apart from strand β7 in the C terminus, which also exhibits high B factors and is disordered in many T1 RNases.

### CSEP0064/BEC1054 interacts with RNA

Due to the structural similarity between CSEP0064/BEC1054 and other RNA-binding proteins, we tested potential association of the protein with RNA and DNA. Firstly, CSEP0064/BEC1054 was labelled with the fluorophore NT-927 and titrated against increasing concentrations of total RNA extracted from barley up to a final concentration of 10 μg/µI in microscale thermophoresis (MST) experiments (Figure 5). An interaction with total RNA was observed, and whilst the complex nature of the substrate precludes accurate calculation of a K_D_, we estimate this to be in the low micromolar range based on an averaged molecular weight for the most abundant RNA species (rRNA) within a pool of extracted total RNA.

**Figure 5.**
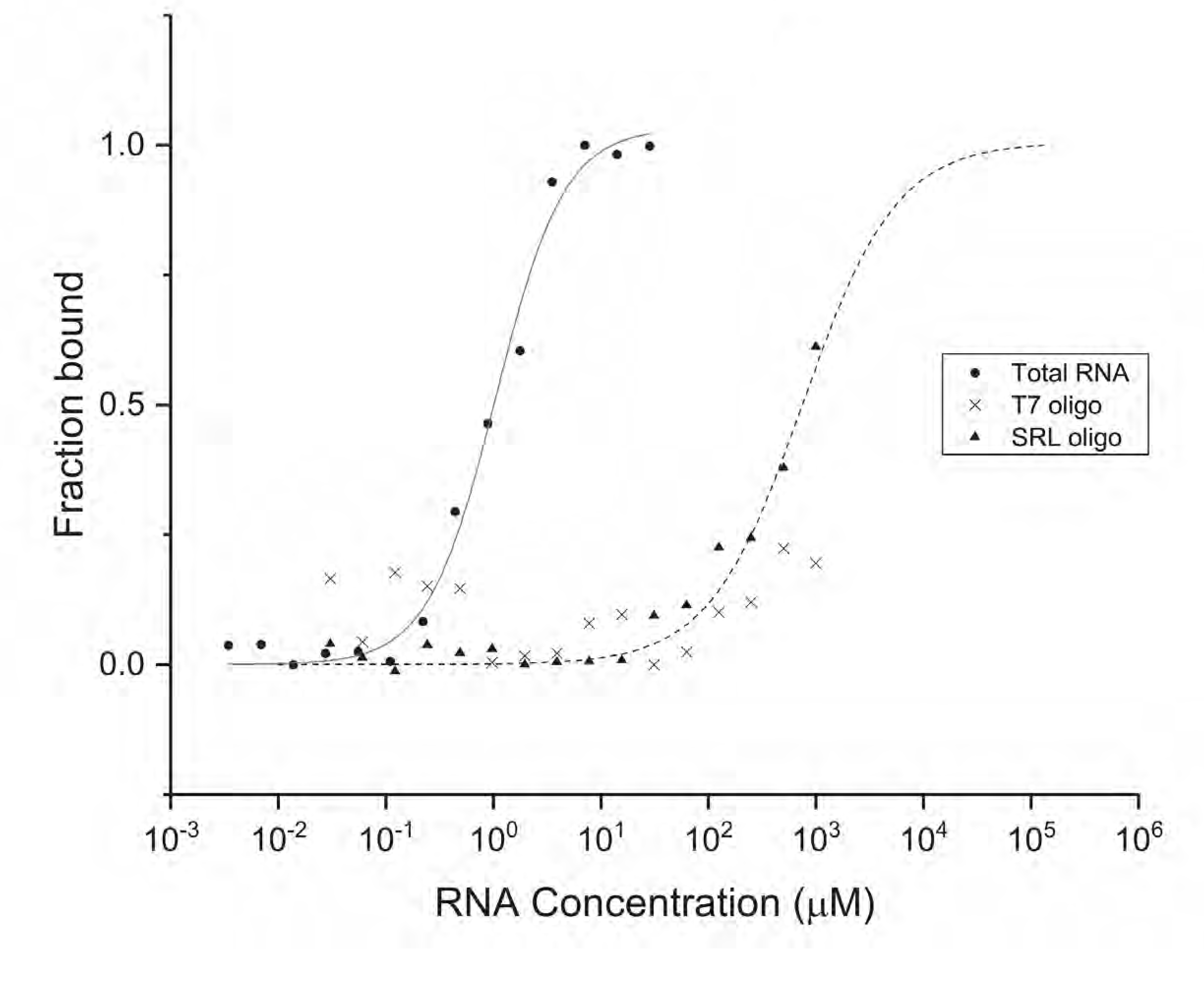
CSEP0064/BEC1054 binds total RNA. Isotherms from microscale thermophoresis assays to determine the *in vitro* RNA-binding capacity of recombinant CSEP0064/BEC1054 protein. Tested RNA species were barley total RNA (•, K_D_ = 1.0±0.1 μM), a ribosomal SRL RNA sequence (▲, K_D_ = 675±52 μM) and a bacteriophage T7 RNA promoter sequence (X). Curves shown are the average of two experimental repeats. In the case of the bacteriophage T7 RNA promoter sequence (X), no curve could be fitted.

To further probe interactions with RNA, the protein was titrated with a specific 32 bp sequence derived from the sarcin-ricin loop (SRL) of the barley 28S rRNA. Affinity for this substrate was over two orders of magnitude weaker than for barley total RNA, with a predicted affinity (K_D_) in the high µM-low mM range (~0.7-1.2 mM; Figure 5).The low affinity of the interaction prevented saturation with the ligand, nonetheless the binding isotherm indicated that this sequence interacts specifically with CSEP0064/BEC1054. In comparison, no binding was observed to an RNA oligonucleotide with the sequence of the bacteriophage T7 promoter, selected as a non-ribosomal RNA control (Figure 5).

Chemical shift perturbation (CSP) analyses using ^1^H-^15^N HSQC NMR spectra were also carried out to observe the interaction of CSEP0064/BEC1054 with the SRL RNA sequence. Titration with 8 mole equivalents of this ligand induced discrete chemical shift changes in a subset of resonances. Titration of the protein with a single-stranded SRL DNA oligonucleotide induced a subset of small CSPs that showed saturation at ~50 mole equivalents (Figure 6A). The continuous shift in the position of resonances upon titration indicated that binding was weak, with a K_D_ above 10 μM (i.e. in fast chemical exchange in NMR timescales). Additionally, many of the crosspeaks that shift are identical to those observed upon addition the SRL RNA oligonucleotide. In these experiments, solution conditions were carefully controlled after each titration to ensure that the observed CSPs were not an artefact from pH changes. CSPs were mapped onto the crystal structure of CSEP0064/BEC1054, and largely localise to the putative RNA-binding site on the concave face of the β-strand (Figure 6B). Residues showing significant shifts include T37, F38 (β3), I39 (β3-β4 loop), V50 (β4), Y79 (β5-β6 loop) and E96 (β7). Residues T37 and F38 correspond to residues T41 and Y42 that form hydrophobic interaction to nucleotides in the crystal structures of T1 RNase, whilst Y79 could have a role in base stacking with the SRL. R25 also exhibits CSPs upon titration, a residue located in a similar region to residues K110, K111 and K113 in the fungal ribotoxin restrictocin, which are in contact with the SRL (Yang et al., 2001). Small CSPs are also observed on the opposite face of CSEP0064/BEC1054. Distal shifts could result from small conformational changes or a non-specific interaction with the SRL ribo-oligonucleotides.

**Figure 6.**
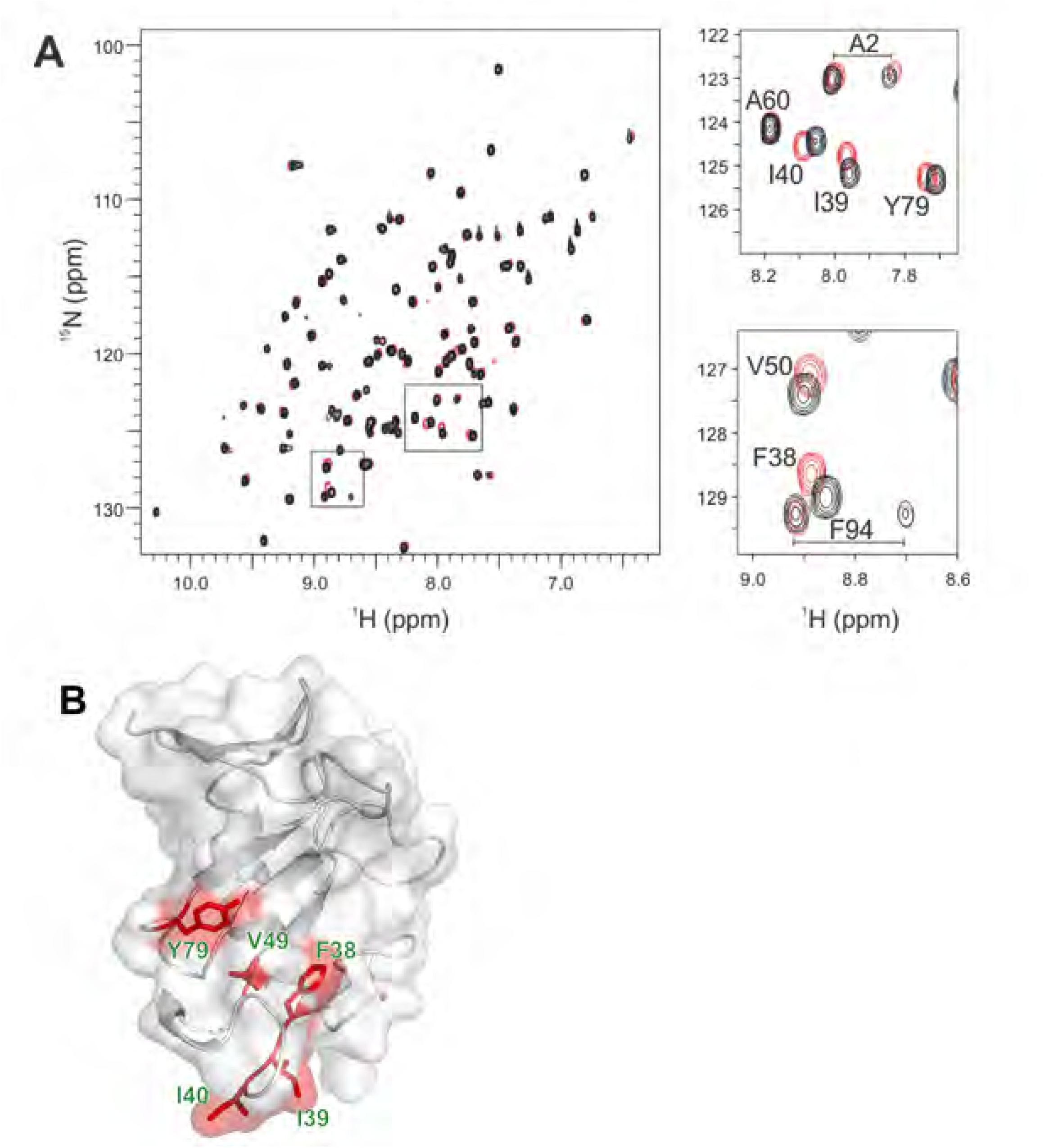
CSEP0064/BEC1054 exhibits discrete chemical shift pertubations that map to its putative RNA-binding site/face. **A**) Overlay of ^1^H-^15^N HSQC NMR spectra of ^5^N-labelled CSEP0064/BEC1054 in the presence (red) and absence (black) of a 50x molar excess of a DNA encoding the sequence of the SRL sequence. Chemical shift pertubations (CSPs) are highlighted in the spectrum and enlarged in separate panels on the right. **B**) Residues that display the most significant CSPs upon ligand binding (as revealed by panel A) mapped onto the crystal structure of CSEP0064/BEC1054.

### CSEP0064/BEC1054 inhibits methyl jasmonate-dependent degradation of host ribosomal RNA

JIP60 is a ribosome-inactivating protein (RIP; (Dunaeva & Goerschen, 1999)). Transcript and protein accumulation of JIP60 is induced by methyl jasmonate (MeJA), a stress-related phytohormone in plants (Reinbothe et al., 1994b). The action of JIP60 on rRNA in plants can be observed by the accumulation of a characteristic degradation product, visible as new RNA species following *in vitro* treatment of total RNA with aniline (Dunaeva et al., 1999). We incubated primary leaves of the transgenic wheat plants constitutively expressing CSEP0064/BEC1054 with MeJA, extracted total RNA, and treated it with aniline. We then analysed the respective RNA fragments by separation in a microchannel-based electrophoretic cell. In the RNA samples obtained from leaves of azygous wheat (negative controls), the aniline treatment resulted in the appearance of a small peak that migrated at an apparent size of about 1,200 bases (Figure 7, Supplemental Figure 2A,B). In the RNA profiles from CSEP0064/BEC1054 transgenic leaves processed in the same way, the area of this peak was reduced. Likewise, this peak was reduced or absent in RNA from CSEP0064/BEC1054 transgenic plant leaves extracted at eight days post inoculation with *B. graminis* f.sp. *tritici* or at five days after MeJa treatment. In the controls with no MeJA induction, the area of the peak was very small or not detectable (Figure 7A). In sum, these data suggest that MeJa treatment triggers fragmentation of rRNA, which is indicated by the occurrence of a presumptive degradation product. This process appeared to be impeded by the presence of CSEP0064/BEC1054 or by powdery mildew infection.

**Figure 7.**
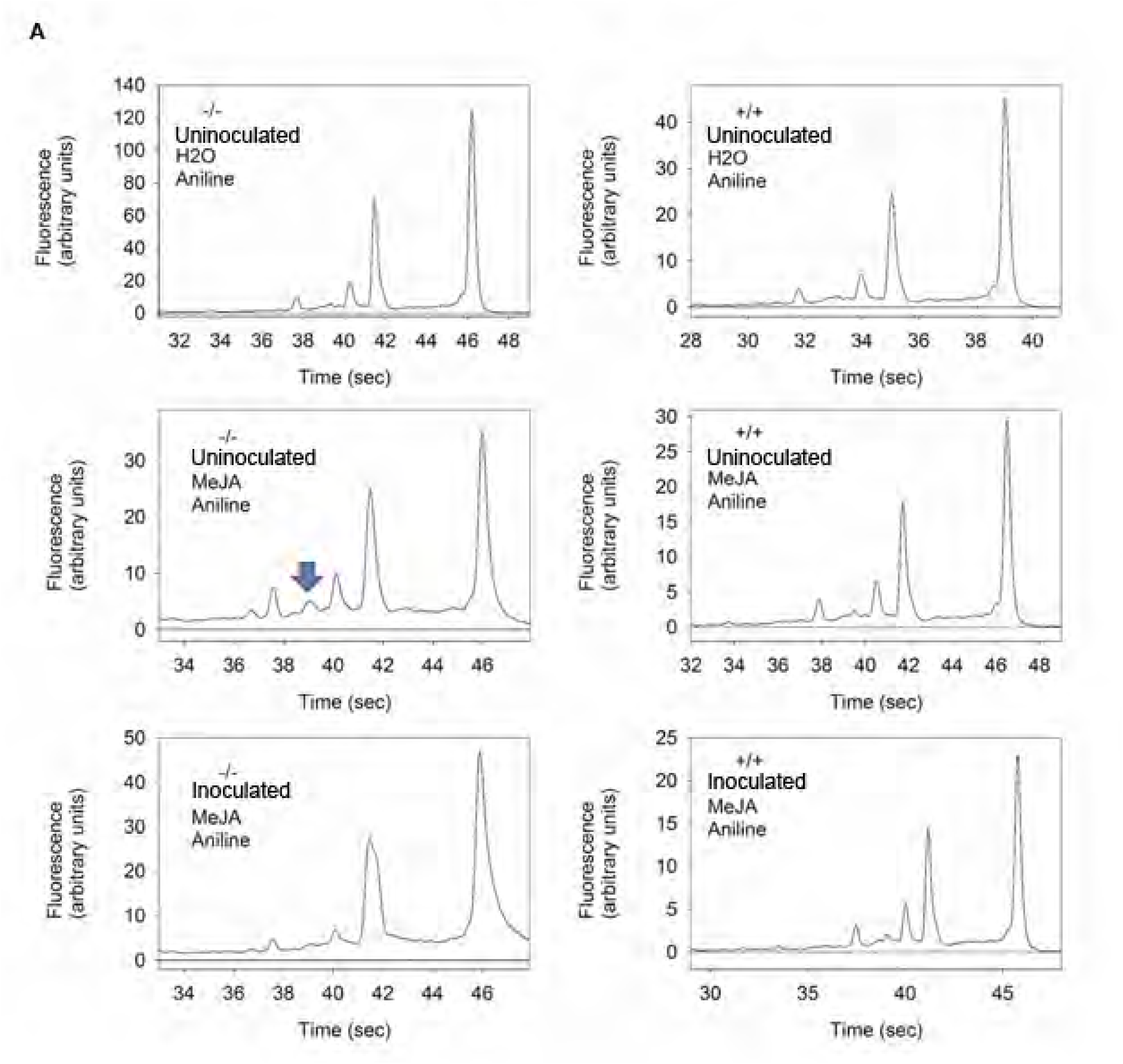
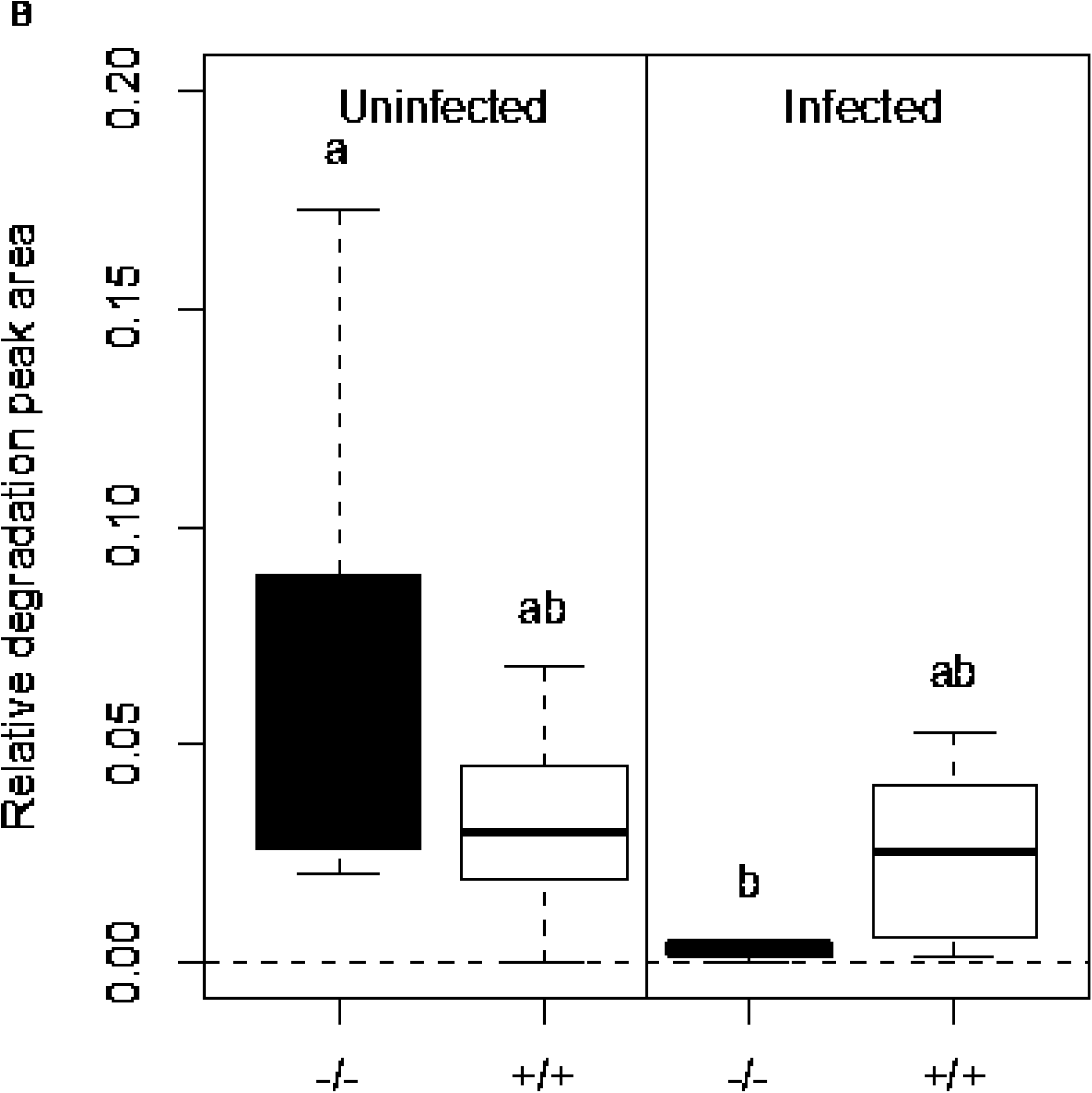
Transgenic expression of CSEP0064/BEC1054 inhibits specific MeJA-dependent aniline-induced rRNA degradation. **A**) Electropherogram peak analysis of total RNA from wheat leaves. Seven day-old transgenic wheat seedlings, either homozygous (+/+) or azygous (-/-) for *wbec1054*, encoding CSEP0064/BEC1054, were inoculated once a day with *B. graminis* f.sp. *tritici* for three days or maintained uninoculated. Primary leaves were harvested, and floated on either water or 40 µM MeJA for a further five days. Following total RNA extraction, RNA was treated with aniline, and then examined using an Agilent Bioanalyzer. The blue arrow indicates the RNA degradation peak of interest, which appears after treatment of the total RNA with aniline. **B**) Quantitative analysis of the assay results shown in **A**. The area of the degradation peak revealed in **A** was calculated in relation to the area of the 28S rRNA of the large ribosomal subunit. *Post-hoc* statistical tests were used to determine whether homozygous (+/+) and azygous (-/-) plants undergoing the same treatment were significantly different, as is indicated by different letters. The boxes represent the quartiles, the thick line denotes the median, and maximum and minimum values are shown by the error bars.

We then measured these effects quantitatively. The area of the peak representing the presumptive rRNA degradation product was determined, relative to the area of the small (18S) and large (28S) rRNAs. The abundance of the newly occurring RNA species was estimated to be up to about 10% of the 28S rRNA peak, depending on the experimental conditions (Figure 7B). The area of the peak of the putative degradation product in RNA from uninfected plants homozygous for the CSEP0064/BEC1054 transgene (line 3.3.14) was lower than in the azygous controls (line 3.3.12). In azygous plants, powdery mildew infection further decreased the formation of the degradation peak, almost completely abolishing it. According to statistical analysis, the infection status had a significant effect on the mean area of the presumptive degradation peak. Overall, powdery mildew infection significantly prevented the formation of the peak. The marked decrease in the degradation peak area also occurred in another independent transgenic line analysed (line 3.3.7; Supplemental Figure 3).

## DISCUSSION

### *In planta* expression of CSEP0064/BEC1054 enhances susceptibility to diverse pathogens

CSEP0064/BEC1054 is a barley powdery mildew candidate effector protein that is highly expressed at early stages of infection (Pennington et al., 2016b). Full expression of the gene is necessary for virulence of *B. graminis* f.sp. *hordei*. (Pliego et al., 2013). Here we investigated further the molecular and structural basis for function of this presumptive effector.

First, we measured the proportion of germinating conidia that lead to the formation of functional haustoria (propH) as deduced by the development of fungal microcolonies on the surfaces of infected leaves on transgenic wheat expressing CSEP0064/BEC1054. PropH was significantly higher in leaves from plants that are homozygous for the transgene, compared to the azygous controls (Figure 1A). We noted differences in the outcome along the leaf blade: the effect of the transgene was more marked at the base of the leaf blades, intermediate in the middle sections and all but disappeared at the apical sections. This is not altogether surprising as leaf maturation in monocots displays a basipetal pattern (Itoh et al., 2005, Wernicke & Milkovits, 1984), and gene expression also occurs in a longitudinally non-uniform manner for leaves of species belonging to the Poaceae (Jiao et al., 2009, Li et al., 2015, Wang et al., 2014). The slope in propH could be related to the basipetal gradients observed for immune-related transcripts. Genes encoding transcriptional regulators, such as WRKY factors, display differential expression along leaf blades, with two genes coding for WRKY proteins being expressed most highly at the leaf tip, and 13 most highly at the leaf base (Li et al., 2015). Several WRKYs play central roles in modulating disease resistance (Pandey & Somssich, 2009). Therefore, a gradient in WRKYs and other proteins that control disease resistance (Li et al., 2015) could explain the differences along the leaf blades in the effectiveness of CSEP0064/BEC1054 in counteracting host defence we observed in our experiments. Similar patterns of susceptibility were also recently observed in barley leaves infected by *P. palmivora* (Le Fevre et al., 2016).

In a separate set of experiments, leaves subjected to *Agrobacterium*–mediated transient expression of CSEP0064/BEC1054 with a C-terminal GFP tag in the dicot *N. benthamiana* were subsequently inoculated with the oomycete *P. tabacina*. Expression of CSEP0064/BEC1054-GFP led to an increase in the *P. tabacina* sporangia formed when compared with a GFP-only control, indicating an increased susceptibility to the oomycete pathogen (Figure 1B).

Taken together, these results point to CSEP0064/BEC1054 affecting plant immunity that limits the infection with pathogens adapted to their specific host. Importantly, the immune pathways targeted by the candidate effector appear to be conserved in monocots and dicots, and affect defence against both fungal and oomycete biotrophic pathogens.

### Interactions of CSEP0064/BEC1054 with host proteins and RNA

Several host proteins interact with CSEP0064/BEC1054 *in vitro* and in yeast (Pennington et al., 2016a). Here, we used BiFC to test whether these host targets associate with CSEP0064/BEC1054 in barley, i.e. in the native cell context of this powdery mildew effector candidate. The fluorescence signals of CSEP0064/BEC1054 and JIP60ml tagged with intact mYFP, as controls, were diffuse and less intense in comparison to the fluorescence of mYFP-tagged candidate interaction partners (Figure 2). Faint signals could be due to poor translation, misfolding and/or rapid turnover of these protein fusions – processes that might be indicative of intolerance of these proteins - at least at high levels - in eukaryotic cells. This scenario would be consistent with interference with cell growth previously observed in yeast expressing CSEP0064/BEC1054 (Pennington et al., 2016a). In the case of JIP60ml-mYFP, this intolerance may be the result of processes leading to cell death induced by its endogenous ribotoxin activity.

Overall, five proteins interacted with CSEP0064/BEC1054 in BiFC experiments in barley: eEF1α(1), eEF1γ, PR5, PR10, and GST (Figure 2B). The fluorescent signals matched the expression and subcellular localization patterns observed for CSEP0064/BEC1054-mYFP and JIP60ml-mYFP. Of these, PR5, GST and eEF1γ interact with CSEP0064/BEC1054 *in vitro* and in yeast, albeit weakly (Pennington et al., 2016a). In Y2H, the tests for PR10 had given inconclusive evidence of interaction due to up-regulation of the β-galactosidase reporter gene, and inconsistent results from the other reporter systems used. The fluorescence observed for eEF1α(1) with CSEP0064/BEC1054 was stronger than for any of the other interactions (Figure 2B). This result contrasts with the previous CSEP0064/BEC1054 protein interaction study, where there was evidence for the interaction of eEF1α(1) *in vitro*, but not in yeast (Pennington et al., 2016a). It is possible that the absence of binding between eEF1α(1) and CSEP0064/BEC1054 in yeast is a false-negative result (Huang & Bader, 2009). If this were the case, it would indicate that other factors or post-translational modification(s) of eEF1α(1) (Dever et al., 1989, Hiatt et al., 1982, L’Italien & Laursen, 1979, Ransom et al., 1998, Toledo & Jerez, 1990) not present in yeast are required for its association with CSEP0064/BEC1054. The specificity of the interaction with eEFα(1) is further underlined by the fact that the pairing CSEP0064/BEC1054-eEFA(3) yielded no detectable BiFC signal.

Overall, the fact that many of these host proteins are capable of consistently associating with CSEP0064/BEC1054 in independent, orthogonal assays (affinity-LCMS, Y2H and BiFC) supports the notion that these interactions may reflect the situation in the true powdery mildew (barley – *B. graminis* interaction) context. The future challenge will be to determine the biological relevance of these events in plant immunity and/or fungal pathogenesis.

The significance of interactions between pathogen effectors and PR proteins may be that they are capable of modulating the antimicrobial activity of these defence-related polypeptides. The PR5 protein identified from barley tissue is a thaumatin-like PR protein (Pennington et al., 2016a). The specific function of PR5 protein(s) is not yet well understood, but it may include antimicrobial and antifungal activity (Hejgaard et al., 1991), and they accumulate in *B. graminis* f.sp. *hordei-*infected barley leaves (Bryngelsson & Green, 1989). An unrelated barley PR protein (PR17c) that is required for defence against *B. graminis* f.sp. *hordei* also physically interacts with a *B. graminis* f.sp. *hordei* candidate effector protein (CSEP0055; (Zhang et al., 2012)). These putative effectors may somehow modulate the antifungal activity of the PR proteins. It remains to be seen whether this is indeed the case for the CSEP0064/BEC1054-PR5 interaction.

The affinity of CSEP0064/BEC1054 for GST could indicate that the effector (or an interaction partner thereof) is glutathionylated by this enzyme. It is known that PR10 is post-translationally S-glutathionylated in birch (Koistinen et al., 2002). The fact that there was no association between GST and JIP60ml, used as a negative control, indicates a degree of specificity in the binding to the candidate effector, possibly related to a post-translational modification.

Our studies also aimed to test whether CSEP0064/BEC1054 binds RNA. Binding isotherms using MST show a clear interaction with total RNA, suggesting that the protein can either bind to RNA unspecifically or that it is capable to recognise a distinct sequence in a well populated species, e.g. an rRNA motif. Experiments using oligonucleotides show that CSEP0064/BEC1054 recognises the SRL of rRNA as a specific ligand, relative to a sequence encoding the bacteriophage T7 promoter, used as a negative control (Figure 5). Using ^1^H-^15^N HSQC NMR spectra, we observed discrete chemical shifts changes upon titration of the SRL RNA oligonucleotide, indicating a weak interaction. This implies that, for a biologically relevant association, the protein may require additional binding elements such as those present in an RNA-protein complex like the intact ribosome, or that a different RNA sequence is recognised *in vivo*.

### Structural similarity of CSEP0064/BEC1054 to RNases and RIPs

Structural modelling of CSEPs from the *B. graminis* f.sp. *hordei* genome (Pedersen et al., 2012) suggested that RNase-type folds are considerably overrepresented relative to other protein folds. For example, CSEP0064/BEC1054 (from CSEP family 21) was predicted to exhibit a fold characteristic to members of the T1 RNase family. As previously noted (Pedersen et al., 2012), *B. graminis* f.sp. *hordei* RALPHs lack the canonical catalytic triad of RNases and are thus unlikely to possess RNase activity. Nonetheless, two main structural features indicated that CSEP0064/BEC1054 may be an RNA-binding protein. In fact, (*1*) the distribution of specific positive charges (Figure 3B) and (*2*) the clustering of aromatic side chains in the concave face of the domain (Supplemental Figure 4) are similar to that observed on the surfaces of F1 RNases involved in RNA ligand binding (Langhorst et al., 1999, Vassylyev et al., 1993). However, it appears that these features would not confer CSEP0064/BEC1054 the same substrate specificity observed in case of these RNases. In particular, this RALPH candidate effector lacks the specific arrangement of backbone and sidechain atoms required for guanosine binding and could instead display a preference for other RNA substrates, or even bind nucleic acids non-specifically, as observed in members of the RNase T2 family (Kurihara et al., 1996).

A defined negatively charged patch on the CSEP0064/BEC1054 surface is evident (Figure 3B). This feature contrasts with the widespread occurrence of positively charged patches in RNA-binding proteins, with an average area five times larger than negatively charged ones (Shazman & Mandel-Gutfreund, 2008). In CSEP0064/BEC1054, this region may serve to bind RNA via a counterion, as observed in the interaction of ribosomal protein L11 to rRNA using Mg^2+^ (Xing & Draper, 1995), or to repel the ligand and electrostatically orient it towards the proposed binding site, as demonstrated for the negatively charged patch of the Gp2 inhibitor of *E. coli* RNA polymerase (Sheppard et al., 2011).

Fungal ribotoxins are another class of proteins related to T1 RNases (Lacadena et al., 2007). Submission of CSEP0064/BEC1054 to the Dali server (Holm & Rosenström, 2010) identified fungal ribotoxins as structural homologues of this effector candidate. Ribotoxins display elongated loop regions and a stretch of lysine residues (K111-K113) that are critical for interaction with the SRL bulged G motif (García-Mayoral et al., 2005, Yang et al., 2001). Although CSEP0064/BEC1054 does not display comparable loop regions, it may specifically recognise RNA and/or protein features on the host ribosome to protect ribosomal RNA from the action of RIPs from the host. Many other plant ribotoxins also lack the triple lysine motif of fungal ribotoxins (including JIP60), and contact the SRL via alternative sequences, namely conserved tyrosine and tryptophan residues that form the “N-glycosidase signature” (Fabbrini et al., 2017).

An additional observation is that CSEPs which are predicted to possess an RNase-like fold, also show some conservation of specific amino acid residues within the putative RNA-binding site but not elsewhere (Supplementary Figure 4). This conservation underscores the potential importance of the role for RALPHs’ RNA-binding for their effector function: effector paralogues may recognise the same host target, and might have diverged under strong evolutionary pressures to escape recognition by surveilling host resistance (R) proteins (Spanu, 2017).

Notably, a CSEP with a predicted RNAse-like fold (CSEP0372) has been recently recognized as a CSEP with *MLA* avirulence activity (AVR_a13_) in *B. graminis* f.sp. *hordei*. Perception of CSEP0372 by the cognate barley nucleotide-binding domain and leucine-rich repeat (NLR) R protein MLA13 results in isolate-specific resistance. Among 17 fungal isolates examined, this *CSEP* gene was found to be present in five allelic variants of which two confer virulence and three avirulence to the fungal pathogen. Another avirulence protein reported in this study (AVR_a1_) shows only weak predicted structural similarity with ribonucleases (Lu et al., 2016). Lately, four additional CSEPs with different *MLA* avirulence activities were identified in *B. graminis* f.sp. *hordei* (AVR_a7_, AVR_a9_, AVR_a10_ and AVR_a22_). When analysed on the basis of the CSEP0064/BEC1054 X-ray structure resolved in the present work (Fig. 3), only AVR_a7_ and AVR_a13_ exhibited significant structural similarity with CSEP0064/BEC1054, while no meaningful structural similarities with CSEP0064/BEC1054 were noted in the case of AVR_a1_, AVR_a9_, AVR_a10_ and AVR_a22_. In summary, these data suggest that allelic MLA immune receptors are capable to mount potent immune responses upon recognition of structurally unrelated CSEPs as avirulence determinants (Saur et al., submitted).

### CSEP0064/BEC1054 interferes with induced rRNA cleavage

The structural similarity of CSEP0064/BEC1054 with RNases and ribotoxins raises the possibility that RALPHs like CSEP0064/BEC1054 bind motifs similar to those recognised by RIPs expressed by the plant host. This prompted us to test the effect of CSEP0064/BEC1054 expression on the integrity of host RNAs. In general, MeJA can induce production of RIPs (Reinbothe et al., 1994b), which cleave an adenine base from the large ribosomal subunit RNA sugar-phosphate backbone (Endo et al., 1988a, Funatsu et al., 1991, May et al., 1989). This exposes the phosphodiester bond in the sugar-phosphate backbone, which can then undergo chemical hydrolysis within the cell (Barbieri et al., 1993, Endo et al., 1988a, Endo & Tsurugi, 1988, Endo et al., 1988b). *In vitro*, the process can be reconstituted by treatment of depurinated rRNA with aniline, which cleaves the sugar-phosphate backbone at the site of the modified nucleotide (Peattie, 1979). This results in the formation of two defined rRNA fragments, ca. 3,000 and 400 nucleotides long, respectively. The smaller rRNA fragment has been observed in barley and is an indicator of RIP activity (Dunaeva et al., 1999). Our experiments showed that RNA extracted from MeJA-induced wheat leaves, treated with aniline, contains a new fragment with an apparent size of about 1,200 nucleotides (Figure 7A). This new RNA species is likely to be a product of depurination/cleavage from a large, abundant RNA, such as rRNA, because it only appears after *in vitro* treatment with aniline. At present we do not know its exact identity, but the size of the RNA fragment is consistent with the products of RNA cleavage previously observed in MeJA-induced RIP, such as JIP60 (Dunaeva et al., 1999). Importantly, the area under the peak varies depending on whether the RNA was extracted from leaves of CSEP0064/BEC1054 transgenic plants or the azygous controls. Furthermore, we observed no additional peaks in RNAs from leaves infected by *B. graminis* f.sp. *tritici* (Figure 7A). These results support the hypothesis that RALPH candidate effectors such asCSEP0064/BEC1054 can interfere with the immune response of the host in dependence of RIP activity.

### Is CSEP0064/BEC1054 an inhibitor of host RIPs?

We have observed an increased susceptibility in different plants to filamentous pathogens upon expression of CSEP0064/BEC1054 (Figure 1). This effect is consistent with the activities displayed by secreted effectors dedicated to modulate or inhibit the immune system of the host (Rovenich et al., 2014). Although the precise activity of CSEP0064/BEC1054 has not yet been determined, the observations made so far suggest an RNA-binding function that counteracts the role of endogenous plant RNases like RIPs. Depurination of a specific nucleotide in the SRL RNA by RIPs (Endo et al., 1987) is a mechanism in plants to limit the spread of fungal infections: it impairs binding of the eEF2/GTP complex to the ribosome (Sperti et al., 1975). This, in turn, inhibits protein synthesis and ultimately leads to apoptosis (Endo & Tsurugi, 1987, Olmo et al., 2001). Thus, suppressing the function of RIPs could be a prime target for CSEP0064/BEC1054 and other proteins in the large family of RALPHs encoded in the *B. graminis* genomes. Several lines of experimental evidence are consistent with this hypothesis. Firstly, our sequence and structural analyses show that this protein is a non-catalytic homolog of fungal RNases (Figure 3C). Secondly, our binding data demonstrate that CSEP0064/BEC1054 is capable of recognising host RNA (Figures 5 and 6A). While the interaction measured is weak, we cannot rule out that the binding affinity of CSEP0064/BEC1054 *in vivo* is enhanced by extended contacts with an RNA motif like the SRL and neighbouring proteins on the ribosomal surface. A similar binding mode has been invoked for ribotoxins and RIPs. For example, structural studies show that the fungal ribotoxin restrictocin can form a stable interaction in solution with the SRL RNA (Yang et al., 2001), and docking of the restrictocin-SRL RNA complex into the structure of the ribosome showed proximity of this enzyme to ribosomal proteins L6 and L14 (García-Mayoral et al., 2005). Thirdly, our data also show that CSEP0064/BEC1054 interacts *in planta* with the elongation factors eEF1γ and eEF1α(1) (Figure 2B). Interestingly, these two proteins form part of the eEF1 complex that binds to the ribosomal A site near to the SRL (Andersen et al., 2006, Unbehaun et al., 2007), and eEF1γ can associate with RIPs (Unbehaun et al., 2007). Other examples that RIPs bind ribosomal proteins include the interaction of fungal trichosanthin with P0, P1 and P2, *in vitro* and Y2H assays (Chan et al., 2007, Chan et al., 2001), the association of pokeweed antiviral protein and L3 in yeast (Hudak et al., 1999, Rajamohan et al., 2001), and the proximity of ricin A-chain *in vivo* to ribosomal proteins L9 and ribosomal stalk protein P0 (Vater et al., 1995).

Significantly, the ribosomal stalk binds elongation factors (Uchiumi et al., 1990). Finally, presence of CSEP0064/BEC1054 *in planta* appears to protect a major host RNA species from MeJA-induced cleavage (Figure 7A), supporting the idea that it competes with a RIP like JIP60 for an RNA-binding site whose integrity is critical for ribosomal function. These observations are consistent with the need of an obligate biotrophic pathogen, such as the powdery mildew fungus, to deal with the consequences of MeJA induction at early stages of infection (Duan et al., 2014).

Given the predicted structural similarity of CSEP0064/BEC1054 with more than 100 paralogs in the *B. graminis* f.sp. *hordei* CSEP repository (Pedersen et al., 2012), it can be expected that additional effector candidates of this pathogen play a similar role. The predicted functional redundancy of these CSEPs might be explained by buffering against the loss of individual effector genes in the highly plastic fungal genome and/or the potential sequential delivery of effector variants during pathogenesis to escape detection by the plant immune system (Thordal–Christensen et al., 2018).

## Conclusions

We hypothesise that the combined interaction of CSEP0064/BEC1054 with ribosomal or ribosomal-associated proteins near the cleavage site could serve to protect the target RNA from the activity of plant RIPs (Figure 8). Preventing the degradation of this rRNA would help to preserve the living cell as a food source for the fungus. This is an essential role for a fungus that is an obligate biotrophic pathogen of plants and goes some way to explain why these candidate effectors are such a prominent component of the CSEP complement in grass powdery mildew fungi.

**Figure 8.** Model of the proposed function of RALPH effectors in the interaction beween powdery mildews and host plants. **A**) Plants recognise potential pathogenic microbes by inducing defence responses, such as the production of the jasmonate-induced protein (JIP60) mediated by methyl jasmonate (MeJA). JIP60 is a ribosome-inhibiting protein (RIP) that degrades host ribosomes and triggers cell death. Host cell death is lethal for obligate biotrophic pathogens such as *B. graminis*. **B**) Powdery mildew fungi secrete abundant RNase-like proteins from their haustoria (RALPH) effectors, such as CSEP0064/BEC1054. We hypothesise that RALPHs facilitate maintenance of live host cells by competing with the host suicide mediated by RIPs such as JIP60.

## METHODS

### Plant and mycelial pathogen growth conditions

Barley, *H. vulgare* L. cv. Golden Promise, and wheat, *T. aestivum* L. cv. Cerco, were grown in Levington Seed and Modular Compost Plus Sand FS2 (Everris, Ipswich, UK) in 13 cm square pots. Plants were kept under long-day conditions, 8 h/16 h hours dark/light cycles, at 21 ^o^C and 33% humidity. Seven days after planting, the barley and wheat seedlings were transferred to 216 dm^3^ Perspex boxes, and inoculated with *B. graminis* f.sp. *hordei* strain DH14 (Spanu et al., 2010), or *B. graminis* f.sp. *tritici* strain “Fielder” (Donal O’Sullivan, NIAB, UK) respectively. The Bimolecular Fluorescence Complementation (BiFC) experiments were performed using the same variety of barley, but grown in 10 cm square pots filled with “Einheitserde” compost (Einheitserde, Frondenberg, Germany). Transgenic wheat experiments were performed using *T. aestivum* L. cv. Fielder, which is also susceptible to the strain of *B. graminis* f.sp. *tritici* used here. *N. benthamiana* was grown in Levington Seed and Modular Compost Plus Sand FS2 (Everris, Ipswich, UK), mixed 2:1 with five-millimetre Vermiculite (Sinclair, Lonconshire, UK) in 9 cm square pots, with one plant per pot. Plants were grown under long-day conditions, with eight hours darkness and 16 hours light, at 25 ^o^C and 33% humidity, and watered twice per week. Four weeks post planting, detached *N. benthamiana* leaves were placed on wet blue-roll (VWR, Chicago, USA), and inoculated with *Peronospora tabacina* (Derevnina et al., 2015) sporangia suspended in autoclaved demineralised water.

### Gene amplification and entry vector construction

Three days post inoculation (dpi) of barley seedlings with *B. graminis,* total RNA was extracted using the using the Qiagen RNeasy Mini Kit (Qiagen, Crawley, UK), and the concentration determined using a NanoDrop-1000 spectrophotometer (Thermo Scientific, Wilmington, USA). Complementary DNA (cDNA) synthesis was performed using 3 μg of barley RNA as a template, with the SuperScript^®^ Double-Stranded cDNA Synthesis Kit (Invitrogen, CA, USA). *Jasmonate Induced Protein 60* cDNA (*jip60*) was amplified from this cDNA template, with PCR conditions as previously described (Pennington et al., 2016a). The resulting PCR product was inserted into the entry vector pCR8 (Invitrogen), as described by the manufacturer. The proximal part of the gene, corresponding to amino acids 1-287 (Genbank CCJR010287053), was re-amplified from the plasmid. The removal of residues 163-185 from this domain (ASTLSGGIGSDVVDDDDGDMLR) is required for RIP activity (Chaudhry et al., 1994). Accordingly, this region was replaced with a methionine-leucine (hence “ml”, residues 181 and 182) linker using the Q5^®^ Site-Directed Mutagenesis Kit (NEB, Ipswich, UK), resulting in a product encoding 267 amino acids. This cDNA/protein is referred to here as “JIP60ml”.

The initial amplification of genes coding for barley proteins to test for interaction with CSEP0064/BEC1054 was performed in the same manner as for JIP60ml. Re-amplification of the plant genes and fungal genes from the pCR8 vector was performed with Phusion High-Fidelity DNA Polymerase (Thermo Fisher Scientific, Schwerte, Germany). The PCR products were either purified using the DNA Clean & ConcentratorTM-5 kit (Zymo Research, Freiburg Germany), or using the QIAquick PCR Purification Kit (Qiagen). The PCR products for pCR8 were treated with restriction enzyme *Dpn*1 (New England Biolabs, Herts, UK) to digest the template plasmid. The remainder were inserted into pDONR201 (Invitrogen, Carlsbad, CA) using the Gateway^®^ BP Clonase^®^ Enzyme Mix (Invitrogen). The resulting entry vectors were transformed into chemically competent TOP10 *E. coli* (Invitrogen), and the transformed bacteria selected on Lysogeny Broth (LB) agar plates containing 100 μg/ml spectinomycin (pCR8) or 50 μg/ml kanamycin (pDONR201). Single colonies from the resulting transformants were picked and grown overnight in LB media with the appropriate antibiotic. Their plasmid DNAs were then purified and sequenced to confirm insertion, orientation and absence of unwanted mutations.

### Vectors for expression in barley and *N. benthamiana* through bombardment or agroinfiltration

Gateway LR Clonase enzyme mix (Invitrogen) was used to recombine the Gateway entry vectors with the Gateway expression plasmids to produce the plasmids for bombardment or for agroinfiltration. The resulting plasmids were transformed into chemically competent TOP10 *E. coli*, and grown overnight on LB medium supplemented with ampicillin. Single colonies of *E. coli* were picked, and the respective plasmids sequenced to confirm the insertion, orientation and absence of unwanted mutations.

### Transient expression in barley

Bacteria, containing the plasmids to be used for biolistic transformation of barley primary leaves, were cultured overnight in 200 μl LB media with ampicillin, and purified using the NucleoBond^®^ Xtra Midi kit (Machery-Nagel, Düren, Germany). Gold micro-carriers (Gold powder, spherical, APS 0.8-1.5 µm, 99.96 (Alfa Aesar, Karlsruhe, Germany), were prepared previously described (Schweizer et al., 1999). In summary, they were weighed in 30 mg aliquots, coated with five µl of DNA solution at 100 ng/μl (one µl of RFP plasmid, two µl of bait plasmid and two µl of prey plasmid; or for the positive expression controls, one µl of RFP plasmid and two µl of expression control plasmid, where all DNA solutions were prepared at 100 ng/μl) whilst being vortexed. Following this, 20 μl of 0.1 M spermidine (Sigma-Aldrich, Munich, Germany) and 50 μl of 2.5 M CaCl_2_ were added dropwise whilst still vortexing. The coated micro-carriers were stored in 60 μl of 100% ethanol on ice until use.

Primary leaves were harvested from eight-day-old barley leaves, and their adaxial surfaces bombarded with coated micro-carriers using a PDS-1000/He^TM^ System with a Hepta^TM^ Adaptor as per the manufacturer’s instructions (Bio-Rad, Munich, Germany). A vacuum of 27 inches of mercury was used with 900 psi rupture disks (Bio-Rad).

Following bombardment, barley leaves were maintained on water agar (1% supplemented with 85 μM benzimidazole. Three days after bombardment, the leaves were coated with perfluorodecalin (95%, Sigma-Aldrich) for a minimum of 30 min before imaging. Leaf samples were analysed using a Leica TCS SP8-X laser-scanning microscope (Leica, Wetzlar, Germany), mounted with a 20x 0.75 numerical aperture water-immersion objective. Transformed cells were identified through observation of RFP/RFP fluorescence, which was excited at 561 nm using the 560 nm diode laser. Fluorescence emission was detected between 600 and 640 nm using a photomultiplier tube detector. Monomeric yellow fluorescent protein (mYFP) fluorescence was excited at 514 nm with an argon laser and fluorescence emission was detected between 520 and 560 nm using a HYD detector. Image capture, analysis and editing were performed using Leica SP8 software. Further editing was performed using FIJI software v2.0 (Schindelin et al., 2012).

### Agroinfiltration and transient expression in *N. benthamiana*

*A. tumefaciens* (GV3101) was transformed with the plasmids by electroporation using a MicroPulser Electroporator as per the electroporator manual (BioRad), followed by recovery for two hours in LB media (1 ml), shaking at 28 ^o^C for two hours. The transformed *Agrobacterium* was subsequently plated onto LB media containing 100 μg of spectinomycin for the plasmids pK7FWG2, pB7RWG2, or pK7WGF2; or 50 μg/ml ampicillin for the colocalisation vectors. Transformed colonies were grown for two days, a single *Agrobacterium* colony selected, the presence of the insert checked by PCR, and the colony streaked onto a fresh plate. Following a further two days of growth, the colonies were resuspended in 2 ml of MMA buffer (10 μM MgCl_2_ and 10 μM MES (2-[N-morpholino] ethanesulfonic acid, pH 5.7). The bacteria were centrifuged (5 min, 8000 g), and resuspended in 2 ml MMA buffer (10 mM). The OD600 was measured, and a bacterial suspension created with a final OD_600_ = 0.5 for RFP constructs, or OD_600_ = 0.2 for GFP constructs.

Three to four weeks old *N. benthamiana* plants were selected for agroinfiltration at OD_600_=0.5. The *Agrobacterium* suspensions were infiltrated into the abaxial surface of leaves (leaves three and/or four from the apex). The leaves were harvested from the plant two to four days after infiltration, and maintained on damp absorbent paper in clear plastic boxes, under long day conditions (16 h/8 h light/dark photoperiod at 18 ^o^C). Infiltrated leaves were mounted in water, and analysed using a Leica SP5 resonant inverted confocal microscope. Excitation and emission wavelengths were 543 nm and 588 nm, respectively, for RFP, 488 nm and 680 nm for plastid autofluorescence, and 488 nm and 495 nm for GFP. RFP was excited with an argon laser, and autofluorescence and GFP were excited using a helium-neon laser. Image analysis and processing were performed using Leica LAS X (Leica Microsystems, Milton Keynes, UK) and Fiji software (ImageJ).

### *N. benthamiana* infection assays

Agrobacteria containing the plasmid pK7FWG2/BEC1054 were infiltrated into one half of an *N. benthamiana* leaf, and Agrobacteria with the GFP-only construct into the other. Both *Agrobacterium* strains were infiltrated at an OD_600_= 0.5. *P. tabacina* (Derevnina et al., 2015) was used to inoculate the leaves within two hours of agroinfiltration. At 10 days after inoculation, inoculated leaves were shaken in water (5 ml) and the number of sporangia harvested were measured through counting with a haemocytometer.

### Generation of transgenic wheat lines

*Wobble CSEP0064/BEC1054* (*wbec1054*) is a synthetic gene corresponding to *B. graminis* f.sp. *hordei CSEP0064/BEC1054*, but lacking the N-terminal signal peptide and containing silent “wobble” mutations, minimizing the sequence identity of the gene with the wild-type *Blumeria* gene; at the same time, the codon usage was optimised for expression in wheat and barley (Pliego et al., 2013). The *wbec1054* gene was cloned into pENTRY (Invitrogen) between the attL1 and attL2 sites, and then recombined into the vector pActR1R2-SCV through LR recombination (Invitrogen). such that the *wbec1054* gene would be expressed from the actin promoter *in vivo* (McElroy et al., 1990). The resulting constructs were transformed into electrocompetent *Agrobacterum* strain AgI1 (Hellens et al., 2000).

The *wbec1054* gene was transformed into *T. aestivum* cv. Fielder through *Agrobacterium*-mediated transformation. Immature seeds were collected at 16-20 days post anthesis, and surface-sterilized (Risacher et al., 2009). Isolated embryos were co-cultivated with *Agrobacterium* at 23^°^C for two days in the dark (Ishida et al., 2015). The embryonic axes were removed, and the subsequent tissue culture performed as described previously (Risacher et al., 2009). The copy number of the *nptII* selectable marker gene was analysed via quantitative real-time PCR (qPCR) using the ΔΔCt method. Plants were grown to maturity, and the T1 generation seeds harvested for further analysis.

### Homozygous line development, genotyping and expression confirmation

Seed dormancy was interrupted by incubation for five days at 32 ^o^C (day), and 4 ^o^C at night. Transgenic wheat seeds were sown in two-by-two centimetre propagation tray chambers, under the growth conditions listed above for wheat and barley, but with the addition of 2 g/l Osmocote Patterned Release Fertiliser (Everris, Ipswich, UK). Seedlings were transferred after two weeks to square plastic pots (9 cm) containing the same potting mixture.

Transgenic wheat DNA (generations T1 to T3) was extracted using the KAPA3G Plant direct PCR protocol (Kapa Biosystems, ROCHE). Transgene presence and copy number was determined for the generations T1 to T4 by qPCR. Analysis was performed through the ΔΔCt method (ΔΔCt=ΔCt _(control gene)_ – ΔCt _(gene of interest)_), with wheat β-tubulin (*tubb6*; U76897.1) as the control, and *wbec1054* as the gene of interest. An expression of 0.00 relative to *tubb6* corresponded to homozygous null plants, referred to hereafter as azygous (-/-), 0.2-0.5 as heterozygous plants (+/-), and 0.5-1 as homozygous plants (+/+).

### Phenotyping transgenic wheat lines

The phenotypic characteristics of mature wheat plants from the T4 generation of lines homozygous (+/+) or azygous (-/-) for *wbec1054* were investigated. Eleven characteristics were assayed: leaf number, maximum height, peduncle (internode 1) and other internode lengths, ear length, subcrown length, fertile tiller number, tiller mass and grain number. Statistical analyses were performed for all characteristics except the subcrown, as the majority of the tiller subcrowns became detached during the drying out phase. The subcrown belonging to the primary tiller could not therefore always be accurately determined.

### Powdery mildew infection assay

Primary leaves were harvested from transgenic wheat plants from the T4 and T5 generations of plants homozygous or azygous for *wbec1054*, and segments (2 cm each) taken from the base, middle and tip using a flat blade. The “mature” leaves correspond to leaf four on 11 week-old plants, “young” leaves were the primary leaves from three week-old plants. The leaf segments were placed on wet blue-roll paper, and inoculated with *B. graminis* f.sp. *tritici* (isolate “Fielder”) on the adaxial leaf surface. One hour post inoculation, leaf segments were transferred onto water agar (0.5% agar supplemented with 16 mg/l benzimidazole) with the infected adaxial side up. Plates were maintained for three days under the growing conditions described above (Plant and mycelial pathogen growth conditions).

Staining of infected leaf segments was performed using 0.1% trypan blue dye in ethanolic lactophenol (1:3.35) (RAL Diagnostics, Martillac, France) for two hours at 80 ^o^C. Destaining was then performed using chloral hydrate (2 mg/ml) for two hours. A Carl Zeiss Axioskop 2 plus microscope (Zeiss, Cambridge, UK) was used to view fungal structures. The proportion of germinated conidia that formed at least one haustorium (propH) was calculated (where propH= (haustorial forming germinated conidia/total number of germinated conidia)). Colonies that formed epiphytic hyphae were used as a proxy for the presence of at least one functional haustorium.

### RNA extraction and analysis

Total RNA was extracted using the Qiagen RNeasy Mini Kit (Qiagen). RNA was quantified with a NanoDrop-1000 spectrophotometer (Thermo Scientific). Aniline (1.2 μl, 1 M)(≥99.5%, Sigma) was used to treat 10 μl, containing up to one µg of extracted RNA in RNase free water, and incubated in the dark at 60 ^o^C for three minutes, following which two µl of 5 M ammonium acetate stop solution with 100 mM EDTA was added and the mixture transferred on ice. The RNA was then precipitated by adding ethanol (1 ml), incubated at −80°C for 20 min, and then collected by centrifugation. The quantity of RNA recovered was measured (NanoDrop 1000) and then analysed using an Bioanalyzer RNA Nano 6000 kit (Bioanalyzer 2100, Agilent Technologies, Santa Clara, CA). The peaks of interest were indentified manually, and the areas under the peaks measured using the manucfacturer’s software (Agilent Technologies 2100 Expert, 2009). The order of the cytoplasmic and chloroplastic ribosomal peaks was obtained from the manufacturer’s guide book (http://citeseerx.ist.psu.edu/viewdoc/download?doi=10.1.1.493.5004&rep=rep1&type=pdf). The sizes of the small and large peaks were obtained from the following PDB models DOI: 10.2210/pdb4v7e/pdb and DOI: 10.2210/pdb5mmj/pdb.

### Production of recombinant CSEP0064/BEC1054 for crystallisation

A gene fusion coding for thioredoxin, a 6xhis tag, a TEV digestion site and the mature form of CSEP0064/BEC1054 (Uniprot N1JJ94, residues 22-118) was expressed in the pNIC-Trx plasmid (kindly provided by Oxford SGC) using the Shuffle T7 Express (NEB) *E. coli* strain. Cell pellets were resuspended in 50 mM Tris, 300 mM NaCl, pH 8.0 (buffer A), and lysed at 25 kpsi using a cell disruptor (Constant Systems Ltd, Warwickshire, UK). Post clarification, supernatants were loaded onto Ni-NTA resin (Qiagen), and washed with buffer A containing 10 mM imidazole prior to elution with buffer A containing 300 mM imidazole. After dialysis in buffer A, CSEP0064/BEC1054 was digested with TEV protease in the same buffer and passed down Ni-NTA resin as described previously to remove thioredoxin and the 6x-his tag, prior to size exclusion chromatography on an S75 16/60 column (GE Healthcare, Buckinghamshire, UK) in 20 mM phosphate pH 7.0, 150 mM NaCl (buffer B).

### Crystallisation of CSEP0064/BEC1054 and structure solution

Purified CSEP0064/BEC1054 was dialysed into crystallisation buffer (10 mM Tris, 150 mM NaCl, pH 7.0) and concentrated to 10 mg/ml for crystallisation. A variety of commercially available solution conditions for crystallisation (Hampton Research,CA, USA) were screened using the sitting-drop vapour diffusion method. CSEP0064/BEC1054 was combined with the mother liquor on a 1:1 ratio in 200 nl drops. Crystals obtained in 0.1 M sodium acetate buffer pH 5.0, supplemented with 30% PEG 4000, 0.4 M (NH_4_)_2_SO_4_ were cryoprotected with 30% glycerol and flash frozen for data collection. A native dataset to 1.3 Å resolution was collected at a wavelength of 0.92 Å using the I04 beamline (DIAMOND Light Source, Oxford, UK) and a second derivative dataset collected at a wavelength of 0.95 Å (above the iodide f’’ edge) from crystals that had been soaked in mother liquor supplemented with 0.5 M sodium iodide. Initial processing, scaling and structure factor calculation of native and iodide SAD datasets was performed using XDS (Evans, 2006, Kabsch, 2010) and TRUNCATE (French & Wilson, 1978) respectively, from within the XIA2 program (Winter, 2010). Phasing of the derivative dataset was performed via single-wavelength anomalous diffraction with density modification, using autoSHARP software (Global Phasing Ltd., Cambridgeshire, UK) (Vonrhein et al., 2011). Following calculation of protein phases, a partial model was built automatically in ARP/wARP (Langer et al., 2008). The model was extended through manual model-buiding in Coot (Emsley et al., 2010)

The refined CSEP0064/BEC1054 structure was thus used as a search model to phase the higher resolution native dataset via molecular replacement using Phaser MR (McCoy et al., 2007). Iterative cycles of model building and reciprocal space refinement were performed in Coot (Emsley et al., 2010) and Refmac5 (Murshudov et al., 1997), respectively until convergence of R_work_ values, where 5% of reflections were excluded for cross-validation.

Both models obtained from SAD and native datasets contained a single copy of CSEP0064/BEC1054 in the asymmetric unit, and all residues were built with the exception of the N-terminal alanine, the side chain of R76 and the C-terminal G97, due to missing or ambiguous density in maps. Model validation was performed using tools in Molprobity (Chen, Arendell et al., 2010).

### CSEP0064/BEC1054 backbone assignment

CSEP0064/BEC1054 was concentrated to 300 μM in buffer B for NMR experiments, where 10% ^2^H_2_O v/v was added to provide a deuterium lock signal. 2D ^1^H^15^N HSQC spectra (Kay et al., 1992, 1994) were recorded at 308 K on a Bruker 600 MHz AvanceIII spectrometer equipped with a TCI cryoprobe (Cross Faculty Centre for NMR, Imperial College London, UK). The chemical shifts of the Cα, Cβ, HN, and CO atoms of the ^13^C, ^15^N labelled CSEP0064/BEC1054 protein were obtained from HNCACB/CBCA(CO)NH and HNCO/HN(CA)CO experiments using standard methods. Data were processed using NMR-Pipe (Delaglio et al., 1995), and analysed in CCPN Analysis (Vranken et al., 2005) or using an in-house version of NMRView (One Moon Scientific) (Marchant et al., 2008).

### BEC1054 RNA binding assays

For microscale thermophoresis experiments, the recombinant CSEP0064/BEC1054 protein was labelled using the Monolith NT.115 protein labelling kit (NanoTemper technologies, Munich Germany) using red fluorescent dye NT-647 NHS (amine-reactive) according to the manufacturer’s instructions. Assays were performed using a Monolith NT.115 MST machine (Nanotemper Technologies), where LED power was kept at 20% and MST power at 40%. Assays were performed in standard or hydrophilic capillaries in 20 mM Tris buffer pH 7.4, 150 mM NaCl, 0.05% Tween, where RNA was titrated against labelled CSEP0064/BEC1054 kept at a concentration of 50 nM. CSEP0064/BEC1054 was titrated with total RNA (extracted from barley) from a starting concentration of 10 μg/ul, or *in vitro* transcribed SRL RNA (5´-ACCUGCUCAGUACGAGAGGAACCGCAGGU-3´) or bacteriophage T7 promoter sequences (5´-AATTTAATACGACTCACTATAGG-3´) from a starting concentration of 1 mM. Curve fitting was performed using the NTAnalysis software (Nanotemper Technologies) in the Thermophoresis + T-Jump mode for the SRL ligand, and using the Hill equation for the total RNA. K_D_ values calculated using non-linear regression.

For NMR experiments, CSEP0064/BEC1054 protein at a concentration of 50 µM was titrated with from 0.5 to up to 80 M equivalents of a DNA SRL oligonucleotide sequence,where 2D ^1^H^15^N HSQC spectra were recorded for each titration point as outlined. Experiments were performed in 50 mM Tris, 150 mM NaCl, at 308 K. Analysis of chemical shift pertubations was performed in CCPN Analysis (Vranken et al., 2005).

### Statistical analyses

Bartlett tests were performed for numerical datasets to determine whether the variance was homogeneous (Crawley, 2005). General Linear Models (GLMs) were conducted on all datasets except for the infection of transgenic wheat with *B. graminis* f.sp. *tritici*, where a Generalised Linear Mixed Model (GLMM) was utilized. Where possible, model simplification was performed, and non-significant or non-interacting factors removed, resulting in the minimal model. For the GLMs, linear hypotheses were tested in a pairwise manner using Games-Howell *post-hoc* tests. The only assay in which a two-way interaction of factors was detected was the RNA extraction and analysis (described above).

A GLMM was used, as non-normal repeated measures (i.e. the fact that the old and young leaves were from the same plants, and the different leaf sections used originated from the same leaves), can be accounted for through the addition of random effects (Crawley, 2005). For the GLMM, the count data corresponding to the total number of germinated conidia (with and without functional haustoria) was bound as a single vector, creating the response variable “y”. To take into account pseudoreplication (due to sampling repeatedly from the same plants/leaves), age, genotype and leaf segment were set as fixed effects, and a binomial family was used due to the response variable being count data. The linear hypotheses were investigated in a pairwise manner using the “multcomp” package in R.

For the phenotyping assay, a “Poisson” family structure was used to account for count data, and datasets were logged to account for overdispersion in the original models (where the residual deviance is much greater than the degrees of freedom).

### Accession numbers

Sequence data from this article can be found in the EMBL/GenBank data libraries under the following accession number(s): *40S 16*, accession KP293844; *eEF1γ*, accession KP293852; *eEF1α(1*) and *eEF1α(3)*, accessions KP293845 and KP293846, *GST*; accession KP293847, *MDH*; accession KP293848, *NDPK*; accession KP293849, *PR10*; accession KP293851, *PR5*; accession KP293851, *JIP60*; accession KY929371 and *CSEP0064/BEC1054* accession CCU83233.1.

## Acknowledgements

This research was funded by the Biotechnology and Biological Sciences Research Council-UK (BB/M000710/1) to PDS, the ERA-CAPS project DURESTrit (PA 861/13) and a project within the Deutsche Forschungsgemeinschaft Priority Programme SPP1819 (PA 861/14-1) to RP

## Supplemental Figure Legends

**Supplemental Figure 1: The phenotype of transgenic wheat is unaffected by the presence of the *wbec1054* transgene, which encodes CSEP0064/BEC1054.**

The T4 generation of transgenic wheat either homozygous (+/+) or azygous (-/-) for CSEP0064/BEC1054 was grown in a random plot design, and the phenotypic characteristics of adult plants were investigated. The boxes represent the quartiles, the thick line denotes the median, and maximum and minimum values are shown by the error bars, and circles indicate outliers. *Post-hoc* tests indicated that none of the phenotypic characteristics were significantly different under the experimental conditions used

Genotyping and qPCR were used to determine that *wbec1054* was present and transcribed in the homozygous wheat line 3.3.14, and absent in the azygous line 3.3.12. Stable transformation leads to the expression of transgenes throughout a plant’s life. It is therefore important to determine whether any changes observed during assays were due to the effect of the transgene on essential processes such as development, or due to the interruption of a gene by the insertion of the transgene (Alberts et al., 2002). *Agrobacterium-*mediated transformation was used to insert the *wbec1054* transgene, and the insertion site for the DNA would have been random. The phenotype of adult wheat azygous or homozygous for *wbec1054* was therefore investigated, under the experimental conditions used for further assays. A randomized block design, with six plots, was used (Borrell et al., 1993, Gasperini et al., 2012) with each block representing one wheat plant. All seeds used to grow the plants were of the same age, and had been stored under the same conditions.

Wheat height (determined using the primary tiller), spike length, leaf number, fertile tiller number, the number of grains per ear, and grain yield are commonly assayed characteristics used to assess wheat plants (Borrell et al., 1993, Fellahi et al., 2013, Hays et al., 2007). The characteristics investigated were not found to be significantly different for the homozygous and azygous plants. These results demonstrate that the presence/absence of *wbec1054* did not affect the adult phenotype of wheat under our experimental conditions.

## Supplemental Figure 2

**(A) Determining Bioanalyzer RNA peak area**. An Agilent Bioanalyzer 2100 was used to measure total RNA was run on a Bioanalyzer RNA Nano 6000 chip. The peak areas were calculated using Agilent 2100 Expert software via manual boundary assignment for the peaks of interest. The asterisk symbol “*” for the small diagnostic peak, “16S” for the small chloroplastic rRNA, “18S” for the small cytoplasmic rRNA, “23S” for the large chloroplastic rRNA and “28S” for the large cytoplasmic rRNA, FU” for fluorescence units.

**(B) Determining the size of the novel RNA peak.** An Agilent Bioanalyzer 2100 was used to measure total RNA was run on a Bioanalyzer RNA Nano 6000 chip. An RNA ladder, consisting of single stranded RNA fragments of known size, was used to estimate the size of peaks within samples of total RNA. The RNA ladder peaks are represented by white circles. The blue cross represents the novel peak, which ran at an approximate size of 1,200 bases. The blue triangles represent the running times and sizes of the chloroplastic and cytoplasmic rRNAs, and are labelled with both their names and sizes, with the abbreviations “16S” for the small chloroplastic rRNA, “18S” for the small cytoplasmic rRNA, “23S” for the large chloroplastic rRNA and “28S” for the large cytoplasmic rRNA.

## Supplemental Figure 3

**Expression of CSEP0064/BEC1054 inhibits specific rRNA degradation induced by MeJA.**

Electropherogram peak analysis of total RNA from wheat leaves (line 3.3.7). The area of the degradation peak was calculated in relation to the area of the large rRNA (28S). Total rRNA was extracted from transgenic wheat plants that were firstly, in case of the infected plants, inoculated with *B. graminis* f. sp. *tritici* for 3 days, and then treated with 40 µM MeJA for the following 5 days. *Post-hoc* tests were used to determine whether +/+ and −/− plants undergoing the same treatment were significantly different, as is indicated by different letters. The boxes represent the quartiles, the thick line denotes the median, and maximum and minimum values are shown by the error bars.

Electropherogram peak analysis of total RNA from wheat leaves (line 3.3.7). The area of the degradation peak was calculated in relation to the area of the large rRNA (28S). Total rRNA was extracted from transgenic wheat plants that were inoculated with *B. graminis* f. sp. *tritici* for 3 days. Infected leaves were then excised and floated on 40 µM MeJA for following 5 days under the same light / dark cycle and temperature conditions the plants had been grown in. *Post-hoc* tests were used to determine whether +/+ and −/− plants undergoing the same treatment were significantly different, as is indicated by different letters. The boxes represent the quartiles, the thick line denotes the median, and maximum and minimum values are shown by the error bars.

## Supplemental Figure 4

**Sequence conservation between *Blumeria graminis* f.sp. *hordei* CSEPs predicted to have an RNAse fold mapped onto the crystal structure of CSEP0064/BEC1054.**

Sequence conservation was calculated using the ConSurf server (Ashkenazy et al., 2016, Landau et al., 2005), where MSAs were supplied from the ClustalOmega server (Analysis Tool Web Services from the EMBL-EBI.(2013 May 13); Nucleic Acids Research 41 (Web Server issue):W597-600). Highly conserved residues are coloured magenta, through white to cyan for regions of low sequence conservation.

Sequence conservation is highest in the core of the protein and the concave face of the main b-sheet; corresponding to the putative RNA binding site when aligned with other RNAses. Aromatics that are also conserved in the T1 RNAse family are indicated.

**Supplemental Table 1.**
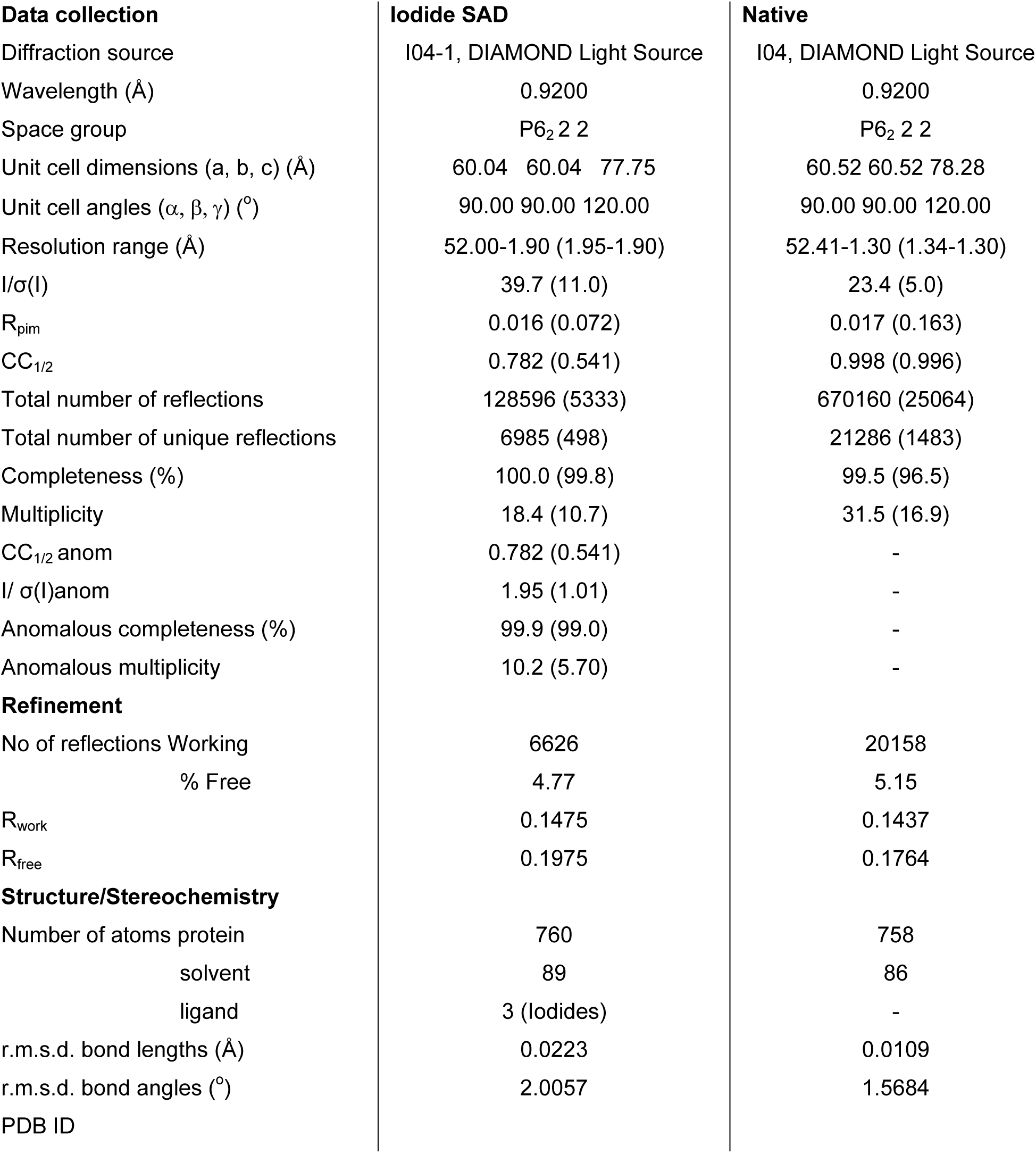
Data processing and refinement statistics for the structures of CSEP0064/BEC0054.

| **Data collection** | **Iodide SAD** | **Native** |
| --- | --- | --- |
| Diffraction source | I04-1, DIAMOND Light Source | I04, DIAMOND Light Source |
| Wavelength (Å) | 0.9200 | 0.9200 |
| Space group | P6<sub>2</sub> 2 2 | P6<sub>2</sub> 2 2 |
| Unit cell dimensions (a, b, c) (Å) | 60.04 60.04 77.75 | 60.52 60.52 78.28 |
| Unit cell angles ($\alpha$, $\beta$, $\gamma$) (°) | 90.00 90.00 120.00 | 90.00 90.00 120.00 |
| Resolution range (Å) | 52.00-1.90 (1.95-1.90) | 52.41-1.30 (1.34-1.30) |
| I/$\sigma$(I) | 39.7 (11.0) | 23.4 (5.0) |
| R<sub>pim</sub> | 0.016 (0.072) | 0.017 (0.163) |
| CC<sub>1/2</sub> | 0.782 (0.541) | 0.998 (0.996) |
| Total number of reflections | 128596 (5333) | 670160 (25064) |
| Total number of unique reflections | 6985 (498) | 21286 (1483) |
| Completeness (%) | 100.0 (99.8) | 99.5 (96.5) |
| Multiplicity | 18.4 (10.7) | 31.5 (16.9) |
| CC<sub>1/2</sub> anom | 0.782 (0.541) | - |
| I/ $\sigma$(I)anom | 1.95 (1.01) | - |
| Anomalous completeness (%) | 99.9 (99.0) | - |
| Anomalous multiplicity | 10.2 (5.70) | - |
| **Refinement** |  |  |
| No of reflections Working | 6626 | 20158 |
| % Free | 4.77 | 5.15 |
| R<sub>work</sub> | 0.1475 | 0.1437 |
| R<sub>free</sub> | 0.1975 | 0.1764 |
| **Structure/Stereochemistry** |  |  |
| Number of atoms protein | 760 | 758 |
| solvent | 89 | 86 |
| ligand | 3 (Iodides) | - |
| r.m.s.d. bond lengths (Å) | 0.0223 | 0.0109 |
| r.m.s.d. bond angles (°) | 2.0057 | 1.5684 |
| PDB ID |  |  |

## ABBREVIATIONS

BiFC: bimolecular fluorescence complementation
CSEP: candidate secreted effector protein
CSP: chemical shift perturbation
eEF: eukaryotic elongation factor
GFP: green fluorescent protein
GO: gene ontology
GST: glutathione S transferase
JIP60: jasmonate-induced protein of 60 kDa
LCMS: liquid chromatography mass spectrometry
MDH: malate dehydrogenase
MeJA: methyl jasmonate
MST: microscale thermophoresis
mYFP: monomeric yellow fluorescent protein
NDPK: nucleoside-diphosphate kinase
NMR: nuclear magnetic resonance
PR: pathogenesis-related
RALPH: RNase-like proteins expressed in haustoria
RFP: red fluorescent protein
RIP: ribosome-inactivating protein
RNase: ribonuclease
rRNA: ribosomal RNA
SRL: sarcin-ricin loop
Y2H: yeast two-hybrid

